# Advancing long-read metagenomic binning via single-copy-gene guided contrastive learning

**DOI:** 10.64898/2026.09.07.749871

**Authors:** Haitao Han, Lauren F. Messer, Christopher Quince, Gary D. Bending, Sébastien Raguideau, Ziye Wang, Shanfeng Zhu

## Abstract

Long-read sequencing advances metagenomics by producing highly contiguous assemblies and more complete metagenome-assembled genomes (MAGs). However, current long-read metagenomic binners fail to incorporate the rich information of long-read assemblies into representation learning and exhibit limited performance on complex datasets. Here, we show that a higher proportion of long-read assembled contigs contain single-copy genes (SCGs) and more SCGs per contig. Therefore, we developed SCGBinner, which leverages SCG-guided contrastive learning to exploit the advantage of long-read data for learning highquality contig embeddings. SCGBinner consistently outperforms other binning methods across five simulated and seven real-world long-read datasets, especially on real-world high-diversity samples. For a deep agricultural soil metagenome, SCGBinner recovered 71% more high-quality MAGs and 38% more near-complete MAGs than the second-best method. Notably, SCGBinner uniquely recovered 449 novel high-quality species, which shed light on the predicted ecological roles of 65 uncharacterised families and 93 novel genera. Overall, SCGBinner could harness the potential of long-read sequencing to provide unprecedented insights into the microbial dark matter of complex microbial communities.

## 1 Introduction

Microbial communities are vital to global biogeochemical cycles and human health [1, 2]. However, most microbes cannot be cultivated in the laboratory. This bottleneck limits our understanding of the microbial world [3]. Metagenomic sequencing has become a widely used culture-independent approach to investigate unknown microbial communities [4]. Typically, sequencing reads are first assembled into longer contigs, followed by binning into metagenome-assembled genomes (MAGs). MAGs have greatly improved our understanding of the microbial tree of life and associated genes and pathways, enabling the discovery of novel metabolisms [5]. Nonetheless, the quality of MAGs directly influences their value [5]. Metagenomic binning is a critical step in recovering MAGs that meet defined quality criteria.

A series of metagenomic binning methods have been developed for short-read sequencing data. For instance, CONCOCT [6], MaxBin 2 [7], MetaBAT2 [8], SolidBin [9], MetaDecoder [10], and MetaBinner [11] use tetranucleotide frequencies (TNF) and coverage information to recover MAGs. Recently, deep learning has been introduced for contig binning. For example, VAMB [12] utilizes a variational autoencoder (VAE) to integrate TNF and coverage information before clustering. SemiBin [13] and COMEBin [14] employ contrastive learning to learn high-quality embeddings for clustering. However, MAGs recovered from short-read metagenomes are often highly fragmented and missing repetitive regions such as the 16S rRNA genes [15].

Long-read sequencing technologies, such as Oxford Nanopore (ONT) and PacBio HiFi, can resolve assembly ambiguities in individual metagenomic samples, thereby enhancing assembly contiguity [16]. The resulting highly contiguous assemblies improve MAG quality, which facilitates more reliable analyses of missing genes and pathways, enables linkage with in situ visualization techniques, and supports the discovery of large operons such as biosynthetic gene clusters [17]. Although binners designed for short-reads can be applied to long-read sequencing data, their performance is not always satisfactory [18]. Therefore, several binning methods have been proposed for long-read sequencing data. HmBin [19] builds on the hifiasm-meta [20] assembler but has been reported to be suboptimal compared to SemiBin [13]. SemiBin2 [21] utilizes a reclustering strategy for short-read data and ensemble-based DBSCAN for long-read data to ensure robust performance on both data types. However, it applies a uniform training process to generate contig embeddings for both shortand long-read data, without considering the different characteristics of longread data. More recently, LorBin [22] employs a two-stage clustering framework with evaluation decision models to enhance MAG recovery, but it follows the short-read binner VAMB [12] to generate contig embeddings using a VAE, similarly overlooking the differences between short- and long-read data.

Whether designed for short or long-read data, there are three components in state-of-the-art binning methods that contribute to improved performance [18]. First, single-copy genes (SCGs) offer valuable information that facilitates contig binning. For instance, MaxBin 2 [7] utilizes SCGs to determine the number of clusters, while MetaDecoder [10] employs SCGs to compute prior probabilities. In addition, MetaBinner [11] and COMEBin [14] leverage SCGs for clustering initialization. Moreover, SemiBin [13] uses SCGs to determine whether the output bins need to be reclustered. Second, contrastive learning is a powerful technique that can generate high-quality representations without labels [23]. For instance, SemiBin2 [21] utilizes contrastive learning with must-link and cannot-link constraints, demonstrating superior performance on long-read data. COMEBin [14] employs multi-view contrastive learning to achieve excellent performance on short-read data. Third, employing an ensemble strategy across multiple clustering results can generate robust binning results. For instance, MetaBinner [11] employs a two-stage ensemble strategy achieving excellent performance on the binning task of the second CAMI challenge (CAMI II) [24]. SemiBin2 [21] uses an ensemble-based DBSCAN for clustering long-read data to improve the robustness of the binning results.

Given that long-read technologies have substantially improved the quality of metagenomic assemblies, we found that the resulting contiguous contigs contain a higher proportion of single-copy genes (SCGs) with more SCGs per contig. This characteristic enables SCGs to provide valuable guidance for representation learning. However, current long-read metagenomic binners fail to incorporate the rich information of long-read contigs into representation learning and exhibit limited performance on complex real-world datasets. Here, we propose SCGBinner, a metagenomic binning method designed specifically for long-read sequencing data. The key innovation of SCGBinner is the integration of SCG information with contrastive learning (SCG-guided contrastive learning). Contigs containing the same SCG harbor homologous coding regions with high sequence similarity, but they are expected to be assigned to different bins. Meanwhile, contrastive learning generates high-quality representations by pulling positive pairs closer and pushing negative pairs farther apart [23]. Therefore, we designed a novel batch construction strategy that considers contigs sharing the same SCG as hard-negative pairs, incorporating them into a single training batch (SCG batch) to fully leverage SCG information during contrastive learning. Subsequently, a Leiden-based ensemble clustering algorithm is applied to further enhance binning performance.

We conduct extensive benchmarking using CAMISIM-generated and de novo-assembled simulated long-read datasets, as well as seven real-world datasets. Across both types of simulated datasets, SCGBinner recovers the highest number of near-complete MAGs (completeness *>*90%, contamination *<*5%) among the state-of-the-art methods on plant and marine datasets. Across all real-world datasets (including soil, marine, sheep rumen, human gut, anaerobic digester sludge, and activated sludge environments), SCGBinner consistently outperforms other state-of-the-art binning methods for both PacBio HiFi and ONT sequencing data. A similar improvement is also observed on real-world short-read datasets. Notably, the benchmarking results demonstrate that SCGBinner significantly improves MAG recovery from complex, high-diversity environments. For a deep agricultural soil metagenome, we demonstrated that SCGBinner could increase the resolution of microorganisms driving both the retention and loss of bioavailable nutrients essential for plant growth and productivity.

## 2 Results

### 2.1 Overview of SCGBinner

We observed highly contiguous contigs assembled from long-read data showed a higher proportion of contigs containing SCGs and a greater number of SCGs per contig (Fig. 1a, b and Supplementary Table 1). Using this information, we propose SCGBinner, a metagenomic binning method designed for long-read data. An overview of SCGBinner is shown in Fig. 1c-e. First, we construct training batches and perform data augmentation. All contigs within each sample are partitioned into standard batches of 1,024. Subsequently, we build SCG batches for all 107 SCGs [7], with each batch consisting of all contigs containing the corresponding SCG. For each contig, we generate six distinct views: one original and five augmented [14]. The augmented views are produced by splitting each contig into three equal-length fragments and two equal-length fragments [21], respectively. Both standard and SCG batches are used to train the combined encoder. Second, we conduct SCG-guided contrastive learning. For each view of a contig, the average embedding of its other five views serves as the positive pair to facilitate faster convergence [25], while the views of all other contigs within the same batch are treated as negative pairs. In this step, the SCG information in SCG batches provides valuable guidance for contrastive learning. Third, a Leiden-based ensemble clustering is performed to obtain the final non-redundant bin set. The trained combined encoder generates embeddings only for the original view of each contig. SCGBinner feeds the resulting contig embeddings into Leiden [26] clustering across a range of parameter values. We use SCGs to calculate the completeness, contamination, and F1-score of bins [21], which are used to identify the optimal partition. SCGBinner then iteratively selects the bin with the highest F1-score among all clustering results, and removes its contigs from every clustering result. To improve efficiency, only the quality of the affected bins are updated [27]. This process continues until no bins meet the quality criteria. Moreover, the remaining bins are collected from the optimal partition to ensure the recovery of potential MAGs that lack the SCGs used in this study (e.g., eukaryotic MAGs). The final output is a non-redundant set of bins, each exceeding 200 kbp.

**Fig. 1.**
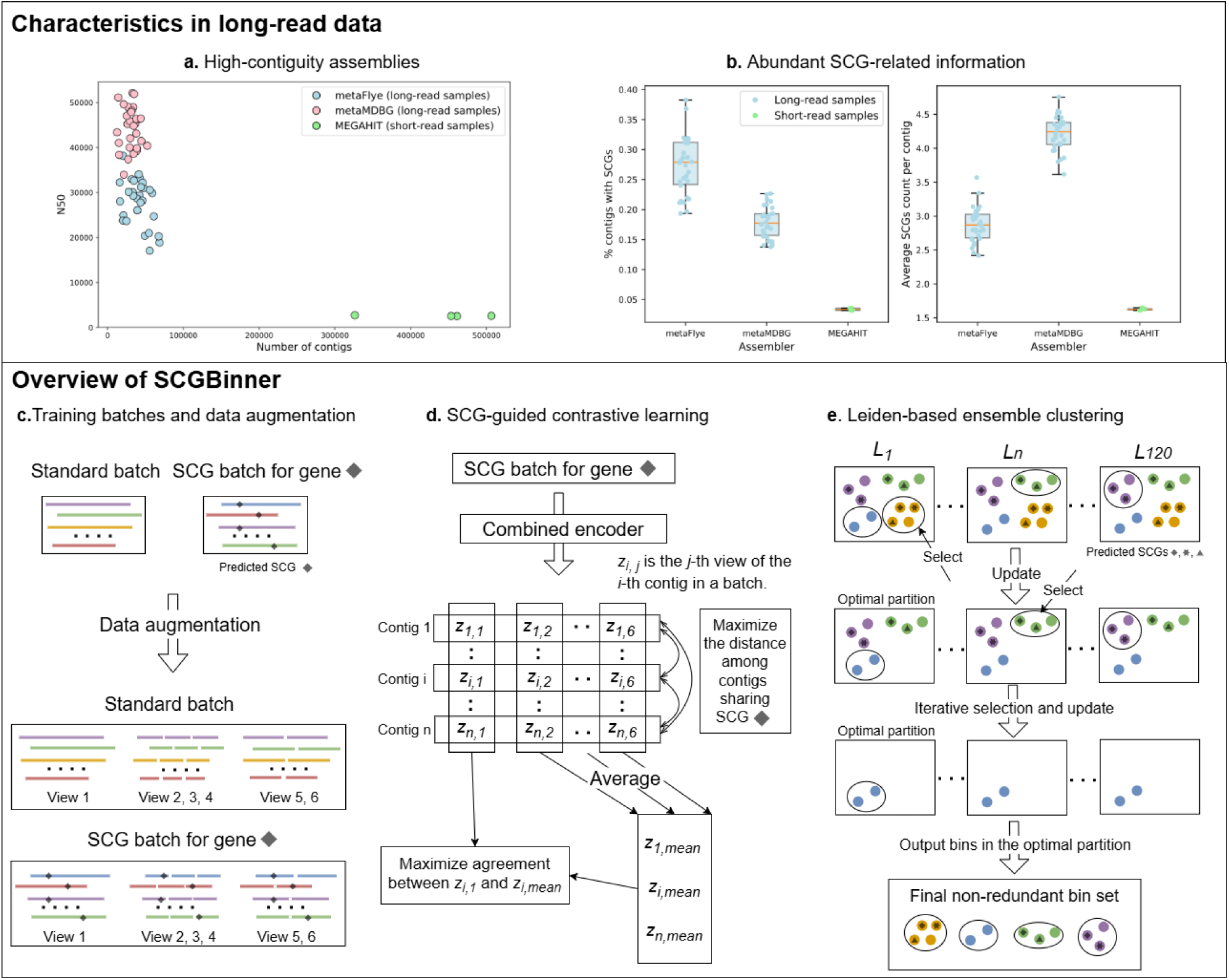
SCGBinner workflow and long-read data characteristics. **a**, N50 and contig counts per sample in the real-world marine dataset (time-series surface seawater metagenomes comprising 30 PacBio HiFi samples and four Illumina samples; see Supplementary Table 14 for details). For 30 long-read samples, assemblies were generated using metaFlye and metaMDBG, while four short-read samples were assembled with MEGAHIT. **b**, Proportion of contigs with SCGs and mean SCGs per contig (calculated among SCG-containing contigs) in the real-world marine dataset. **c**, Training batches and data augmentation: contigs within each sample were partitioned into standard batches of 1,024, and SCG batches were constructed for all 107 SCGs, each comprising all contigs carrying the corresponding SCG. Each contig is segmented into three equal-length fragments and two equal-length fragments, respectively, producing a total of six distinct views: one original and five augmented. **d**, SCG guided contrastive learning: the combined encoder is trained with both standard and SCG batches to generate high-quality embeddings for each contig. **e**, Leiden-based ensemble clustering: generate final nonredundant bin set.

### 2.2 SCGBinner improved the recovery of MAGs on simulated datasets

We compared SCGBinner with other state-of-the-art methods on simulated plant, marine, and strain-madness metagenomic datasets (Fig. 2, Supplementary Figure 1 and Supplementary Table 2). These simulated long-read datasets were generated using CAMISIM [28], based on the reference genomes and genome abundances provided by CAMI II [24] (see Section 4.6). The number of recovered bins, accuracy (bp), and F1 score for sample (bp) were calculated using AMBER (version 2.0.3) [29].

**Fig. 2.**
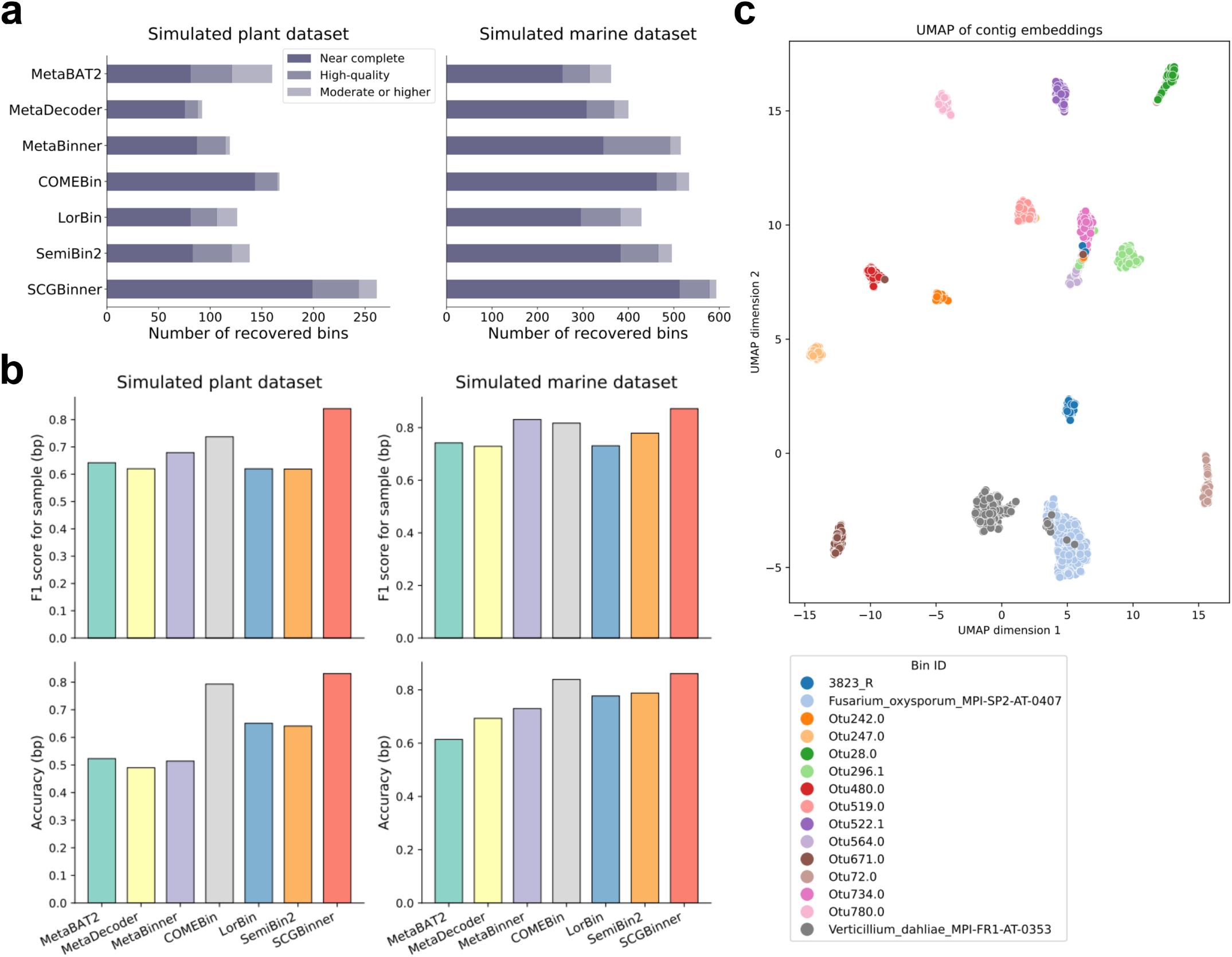
Comparison of binning methods on two simulated long-read datasets generated using CAMISIM: the plant dataset containing 468 complete genomes and the marine dataset containing 775 complete genomes. **a,** The number of recovered near-complete, high-quality, and “moderate or higher” quality MAGs. **b,** F1 score for sample (bp) and accuracy (bp) of binning methods. **c,** UMAP visualization of contig embeddings generated by SCGBinner on the simulated plant dataset (15 genomes with more than 100 contigs).

SCGBinner outperformed all other state-of-the-art methods in MAG recovery on the simulated plant, marine, and strain-madness datasets (Fig. 2 and Supplementary Figure 1). On the plant dataset, SCGBinner recovered 56%, 47%, and 39% more “moderate or higher” quality (completeness *>*50%, contamination *<*10%), high-quality, and near-complete MAGs, respectively, than the second-best method. We also found that SCGBinner recovered 11 “moderate or higher” quality eukaryotic MAGs, whereas the second-best binner, COMEBin, and the widely used SemiBin2 recovered no “moderate or higher” quality eukaryotic MAGs. These results suggest that SCGBinner provides advantages over existing advanced binning methods through recovery of both prokaryotic and eukaryotic genomes on this plant dataset. Moreover, SCGBinner outperformed the second-best binners by 10.3% in F1 score for sample (bp) and 3.8% in accuracy (bp). On the marine dataset, SCGBinner improved MAG recovery by 11%, 14%, and 10% for “moderate or higher” quality (completeness *>*50%, contamination *<*10%), high-quality, and near-complete MAGs, respectively, and improved F1 score for sample (bp) and accuracy (bp) by 4.1% and 2.2%, respectively, compared with the second-best method. SCGBinner also recovered the highest numbers of “moderate or higher” quality, high-quality, and near-complete MAGs on the simulated strainmadness dataset (Supplementary Figure 1), while achieving competitive F1 scores for sample (bp) and accuracy (bp). Furthermore, we randomly selected 15 genomes with more than 100 contigs from the simulated plant dataset and visualized their corresponding contig embeddings using UMAP [30] (Fig. 2c and Supplementary Figure 2). Compared with the contig embeddings generated by other binning tools, the SCG-guided contig embeddings form more compact clusters with clearer separation between clusters (Supplementary Table 4).

In addition to the simulated gold-standard assemblies generated by CAMISIM, we also evaluated the binning methods on simulated datasets that were de novo assembled from long reads generated with Badread [31] (Supplementary Note 2 and Supplementary Figure 3). SCGBinner consistently outperformed the other methods on the simulated marine and plant datasets, yielding the largest numbers of MAGs across the “moderate or higher” quality, high-quality, and near-complete categories. However, the extreme strain diversity in the strain-madness dataset posed substantial challenges for current long-read assemblers (Supplementary Table 3), resulting in poor assembly quality and consequently limiting MAG recovery across all evaluated binning methods.

### 2.3 SCGBinner improved the recovery of MAGs on real-world datasets

We next compared SCGBinner with other methods on seven real-world metagenomic datasets, covering soil [32, 33] (one 250 Gb PacBio HiFi sample and six Oxford Nanopore samples), marine [34] (30 PacBio HiFi samples and four Illumina samples), sheep rumen [35] (one PacBio HiFi sample), anaerobic digester sludge [15] (AD sludge, three PacBio HiFi samples), human gut [15] (four PacBio HiFi samples), and activated sludge [36] (23 Oxford Nanopore samples and 23 Illumina samples) environments. Illumina samples were incorporated to generate the hybrid-assembled datasets (each long-read sample and its corresponding short-read sample were assembled together).

We first benchmarked binners on PacBio HiFi samples assembled with metaFlye (Fig. 3a). SCGBinner substantially outperformed the other binners in recovering a greater number of high-quality and near-complete MAGs across all datasets using single-sample binning (for single-sample binning, each sample is assembled and binned individually; see Section 4.2). In contrast, SemiBin2, LorBin, and MetaBinner showed lower performance. Specifically, SCGBinner recovered 66-114%, 70–103%, 28–59%, 36–48%, and 25–76% more near-complete MAGs than these three methods on the soil, marine, sheep rumen, human gut, and AD sludge datasets, respectively. For multi-sample binning (each sample is assembled and binned individually, but the coverage feature is calculated from all samples; see Section 4.2), SCGBinner retrieved 89%, 79% and 73% more near-complete MAGs than SemiBin2, LorBin and MetaBinner on the marine dataset, respectively (Supplementary Figure 4a). Similar trends were observed on Nanopore samples assembled with metaFlye (Supplementary Figure 5).

**Fig. 3.**
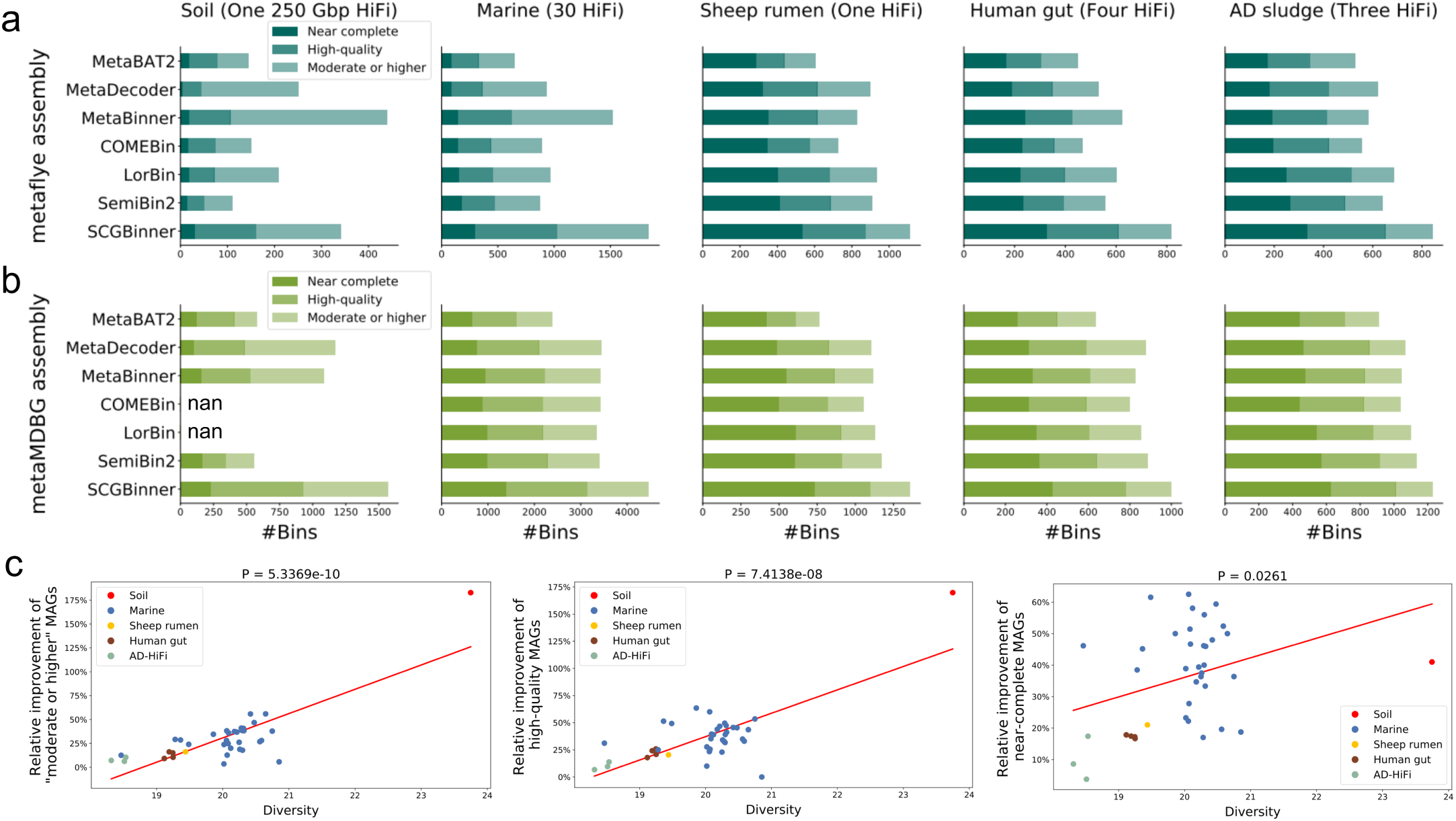
Comparison of binning methods on five real-world long-read datasets using single-sample binning. **a,** The number of near-complete, high-quality, and “moderate or higher” quality MAGs recoverd from contigs assembled by metaFlye. **b,** The number of near-complete, high-quality, and “moderate or higher” quality MAGs recoverd from contigs assembled by metaMDBG. “nan” denotes that the corresponding binners failed to produce binning results. COMEBin failed due to memory-related errors when executed with 16 threads on a machine with 503 GB of RAM, whereas LorBin was unable to complete the binning process on the same machine after one month of execution. **c,** Sample-wise diversity and relative improvements in MAG recovery. We used the Spearman method to compute the correlation coefficient. Sample diversity was estimated using Nonpareil [37].

Then, we benchmarked binners on PacBio HiFi samples assembled with metaMDBG (Fig. 3b). For single-sample binning, SCGBinner consistently recovered substantially more near-complete MAGs than SemiBin2, LorBin, and MetaBinner, with increases of 38–40%, 40–47%, 19–32%, 17–28%, and 10–31% across soil, marine, sheep rumen, human gut, and AD sludge datasets, respectively. For multi-sample binning, SCGBinner retrieved 24%, 23% and 53% more near-complete MAGs than SemiBin2, LorBin and MetaBinner on the marine dataset, respectively (Supplementary Figure 4b). The performance gains of SCGBinner were even more pronounced under the co-assembly binning, with SCGBinner recovering 50–66%, 47–104%, and 18–42% more near-complete MAGs than SemiBin2, LorBin, and MetaBinner across the marine, human gut, and AD sludge datasets, respectively (Supplementary Figure 6). Moreover, all binners recovered higher numbers of “moderate or higher” quality, high-quality, and near-complete MAGs from metaMDBG assemblies than from metaFlye assemblies across all datasets. This is attributable to metaMDBG generating higher-quality assembled contigs that enhance binning performance, while SCGBinner consistently achieves substantial improvements over the other binners on assemblies from both metaFlye and metaMDBG. Following the MIMAG standards [38], we further examined the presence of the 23S, 16S, and 5S rRNA genes and at least 18 tRNAs in near-complete MAGs recovered by all binners across all real-world longread datasets (Supplementary Table 5 and 6). Overall, SCGBinner achieved the best performance in recovering near-complete MAGs containing all three rRNA genes and at least 18 tRNAs.

To examine whether SCGBinner also improves binning performance on shortread metagenomic datasets, we benchmarked binning methods using Illumina samples assembled with MEGAHIT (Supplementary Figure 7). SCGBinner recovered the highest number of “moderate or higher” quality, high-quality, and near-complete MAGs on both the marine and activated sludge datasets. COMEBin achieved the second-best performance. Specifically, SCGBinner recovered 55% and 25% more near-complete MAGs than COMEBin on the marine dataset, using single-sample and multi-sample binning, respectively. On the activated sludge dataset, SCGBinner recovered 24% and 15% more near-complete MAGs than COMEBin using single-sample and multi-sample binning, respectively. Similar trends were observed on hybrid assembled datasets (Supplementary Figure 8).

### 2.4 SCGBinner improved the recovery of MAGs from real-world high-diversity communities

To evaluate the correlation between the performance of SCGBinner and microbial diversity in real-world datasets, we used Nonpareil [37] to estimate the average coverage and diversity of PacBio HiFi samples (Fig. 3c and Supplementary Figure 9). The Nonpareil curves illustrate the sequencing effort required for different communities to reach the same level of coverage, with more diverse communities demanding greater sequencing effort (Supplementary Figure 9). The soil sample exhibited the highest diversity, followed by the marine, sheep rumen, human gut, and AD sludge samples. Consistently, greater improvements in MAG recovery were observed for SCGBinner in higher-diversity samples. For example, compared to SemiBin2, a widely used longread binning method, SCGBinner achieved increases of 40%, 40%, 19%, 17%, and 10% in the number of near-complete MAGs recovered from the soil, marine, sheep rumen, human gut, and AD sludge datasets, respectively (Fig. 3b). Therefore, we evaluated the correlation between diversity and the relative improvement in MAG recovery by SCGBinner compared to SemiBin2 across all PacBio HiFi samples (Fig. 3c). A significant positive correlation was observed (*P* = 5.3369 *×* 10*^−^*^10^ and *R* = 0.8073 for “moderate or higher” quality MAGs; *P* = 7.4138 *×* 10*^−^*^8^ and *R* = 0.7398 for highquality MAGs; *P* = 0.0261 and *R* = 0.3560 for near-complete MAGs), suggesting that SCGBinner improves the recovery of MAGs from high-diversity communities.

### 2.5 Broader microbial diversity in MAGs recovered by SCGBinner

To analyze the microbial diversity of high-quality MAGs (HQ MAGs; contamination *<*10% and completeness *>*70%) recovered from the 250 Gbp soil HiFi sample (deep agricultural soil metagenome) assembled using metaMDBG, all HQ MAGs generated by SCGBinner and two widely used binning methods (SemiBin2 and MetaBAT2) were clustered into 1,042 species-level groups. These species were then taxonomically annotated using the Genome Taxonomy Database toolkit (GTDB-Tk, version 2.4.0) with GTDB release r220, and the corresponding phylogenetic trees were constructed (Fig. 4, Supplementary Figures 10 and 11). MAGs recovered by SCGBinner contributed to large parts of the tree (Fig. 4a). Notably, for known taxa, two phyla, five classes, 22 orders, 43 families, and 84 genera were found only by SCGBinner (Fig. 4b). In comparison, one order, one family, and one genus were specific to SemiBin2, while three orders, five families, and seven genera were specific to MetaBAT2. Moreover, a total of 1,018 HQ species-level MAGs were novel compared to GTDB r220. Among these MAGs, SCGBinner recovered 449 unique novel species, 93 unique novel genera, and five unique novel families, whereas SemiBin2 recovered 33 unique novel species and four unique novel genera, and MetaBAT2 recovered 60 unique novel species, 17 unique novel genera, and one unique novel family (Fig. 4c and Supplementary Figure 11). These results demonstrate the advantage of SCGBinner in resolving complex microbial communities across both known and unknown taxa.

**Fig. 4.**
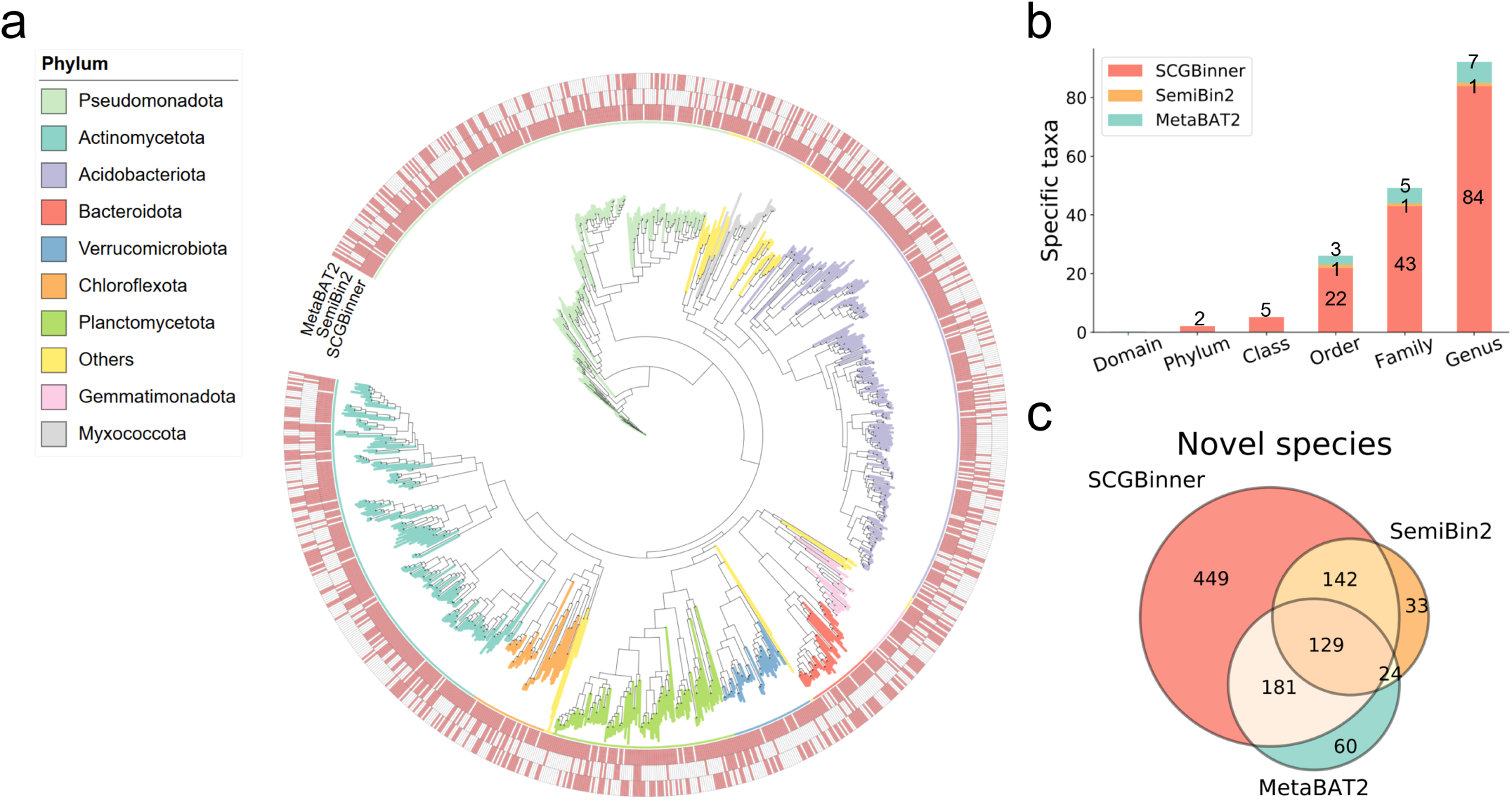
GTDB annotation of HQ species recovered from the deep agricultural soil metagenome). **a,** Phylogenetic tree of species-level HQ MAGs based on GTDB-Tk bacterial marker genes. **b,** Unique taxa per binner at different taxonomic levels. **c,** Number of novel species per binner.

### 2.6 SCGBinner reveals extensive potential ARG hosts and novel BGCs

The recovery of novel species from complex environments provides access to potential untapped reservoirs of biosynthetic gene clusters (BGCs) that produce natural products with potential pharmaceutical value [39]. To identify BGCs in HQ species, MAGs recovered from the deep agricultural soil metagenome by MetaBAT2, SemiBin2, and SCGBinner were independently clustered into HQ species. BGCs within these species were then identified and compared to the 1,225,071 known BGCs in the BiG-FAM [40] database using BiG-SLiCE [41] to assess their novelty. Potential novel BGCs were identified at BiG-SLiCE distance thresholds of 900 (default), 1,200, and 1,500 (Fig. 5a), where higher thresholds correspond to more stringent criteria. HQ species recovered by SCGBinner uncovered 168% and 99% more novel BGCs (threshold = 900) compared to SemiBin2 and MetaBAT2, respectively. Similar results were observed under more stringent thresholds. In total, 49, 42, and 44 BGC types were detected in the HQ species recovered by SCGBinner, SemiBin2, and MetaBAT2, respectively (Supplementary Table 7). Notably, SCGBinner recovered a substantial number of BGCs across a wide range of BGC types encoding chemically diverse secondary metabolites with potential agrochemical, pharmaceutical, and nutraceutical relevance, such as terpenes, T3PKS, NRPS, RiPPs, lasso peptides, and beta lactones (Fig. 5b). The taxonomic origins of BGCs specific to each binner were further analyzed (Fig. 5c, Supplementary Table 8). HQ species recovered by SCGBinner contained 1,627 unique novel BGCs across 18 phyla, representing 5.0- and 13.1-fold increases over MetaBAT2 (328) and SemiBin2 (124), respectively. Moreover, MAGs recovered by SCGBinner expanded the identification of novel BGCs across the Acidobacteriota and Actinomycetota, phyla well-known for their biosynthetic potential in soil. Even within the less well-known phyla, such as the Gemmatimonadota, Methylomirabilota, and Desulfobacterota B, SCGBinner allowed the detection of novel BGCs that would otherwise be overlooked in MAGs recovered using SemiBin2 or MetaBAT2. These results suggest that the application of SCGBinner to long-read metagenomic datasets may facilitate the discovery of novel secondary metabolites across diverse phyla.

**Fig. 5.**
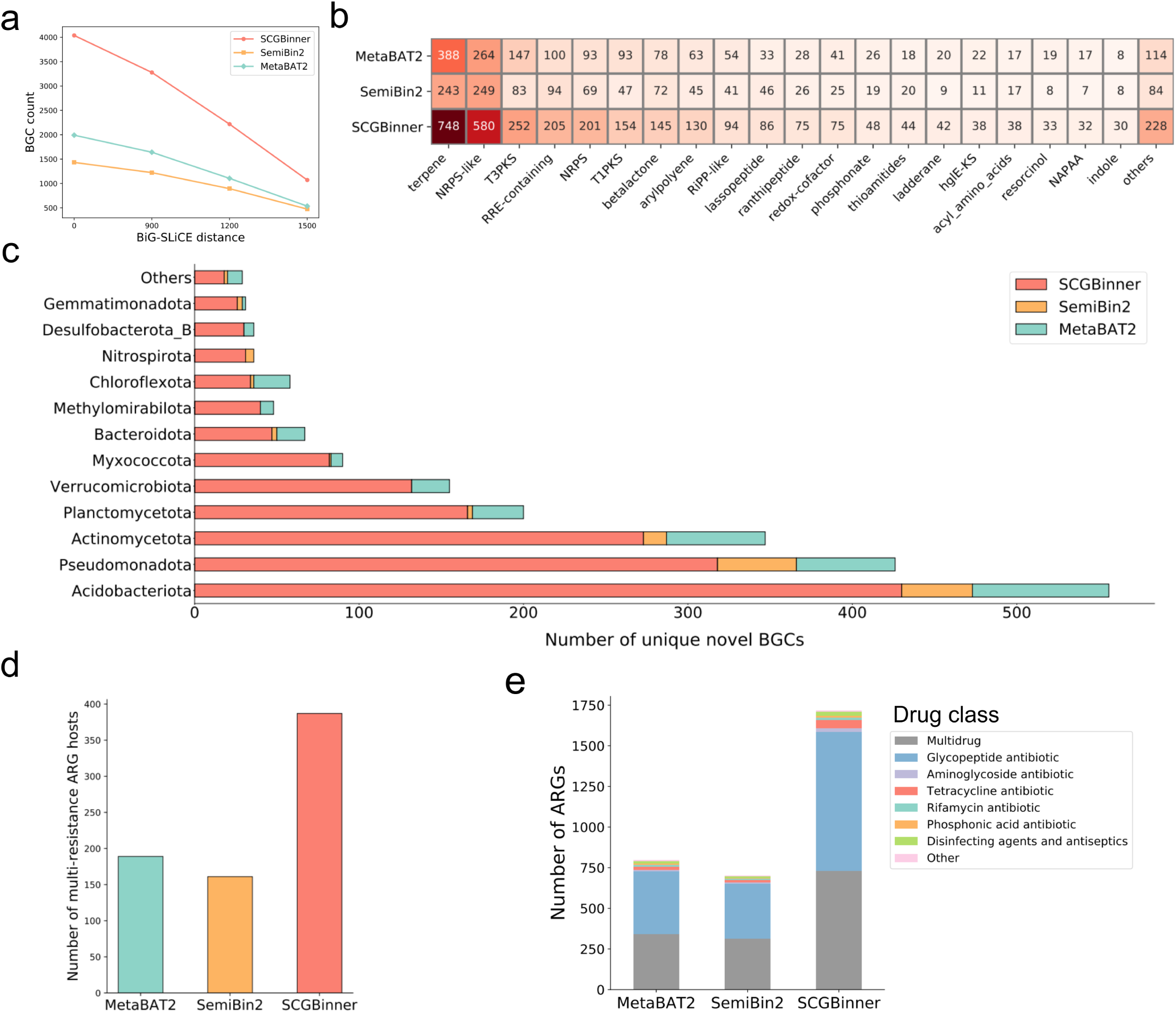
Potential ARGs and BGCs identified in HQ species recovered from a deep agricultural soil metagenome. **a,** Number of BGCs at different BiG-SLiCE distance thresholds. **b,** Number of novel BGCs across different BGC types. **c,** Number of unique novel BGCs across different phyla. **d,** Number of potential multi-resistance ARG hosts. **e,** Number of ARGs in HQ species and drug class distribution.

In addition to biosynthetic genes, soil microbiota are known reservoirs of antibiotic resistance genes (ARGs), which pose a serious threat to public health if transferred to human and animal pathogens [42]. Here we identified potential ARGs within HQ species recovered from the deep agricultural soil metagenome. To ensure reliability, only ARGs located on contigs longer than 10 kb were considered [43]. MAGs carrying more than one ARG type were regarded as potential multi-resistance ARG hosts [44]. Notably, SCGBinner revealed 140% and 104% more potential multi-resistance ARG hosts compared to SemiBin2 and MetaBAT2, respectively (Fig. 5d). Furthermore, HQ species recovered by SCGBinner carried 132% and 113% more ARGs conferring resistance to at least two drug classes (multidrug resistance) than those recovered by SemiBin2 and MetaBAT2, respectively (Fig. 5e). The improved genomic resolution afforded by SCGBinner may facilitate risk assessment and management of antibiotic resistance in complex microbial environments such as agricultural soil.

### 2.7 Functional potential of MAGs recovered by SCGBinner

Metabolic reconstruction of HQ species (Section 2.6) recovered by MetaBAT2, SemiBin2, and SCGBinner from the deep agricultural soil metagenome revealed the functional potential of the agricultural soil microbiota. In total, distinct KO types identified in HQ species recovered by SCGBinner, SemiBin2, and MetaBAT2 were as follows: 9,472; 8,021; and 8,593, respectively. Among these, 780, 27, and 116 KO types were exclusively detected in HQ species recovered by SCGBinner, SemiBin2, and MetaBAT2, respectively. This suggests that while each binner may recover different functional groups, the MAGs recovered by SCGBinner provide a more comprehensive metabolic reconstruction of the microbial community in addition to recovering more MAGs. Indeed, compared with SemiBin2 and MetaBAT2, HQ species recovered by SCGBinner exhibited more diverse functional profiles (Fig. 6a) and more complete key carbon, nitrogen, and sulfur cycling pathway modules. For these pathways, 34, 31, and 32 different types of complete modules were identified in HQ species recovered by SCGBinner, SemiBin2, and MetaBAT2, respectively. In total, these corresponded to 8,620, 3,405, and 4,091 complete modules for the three methods, respectively (Fig. 6b-d and Supplementary Tables 9-11).

**Fig. 6.**
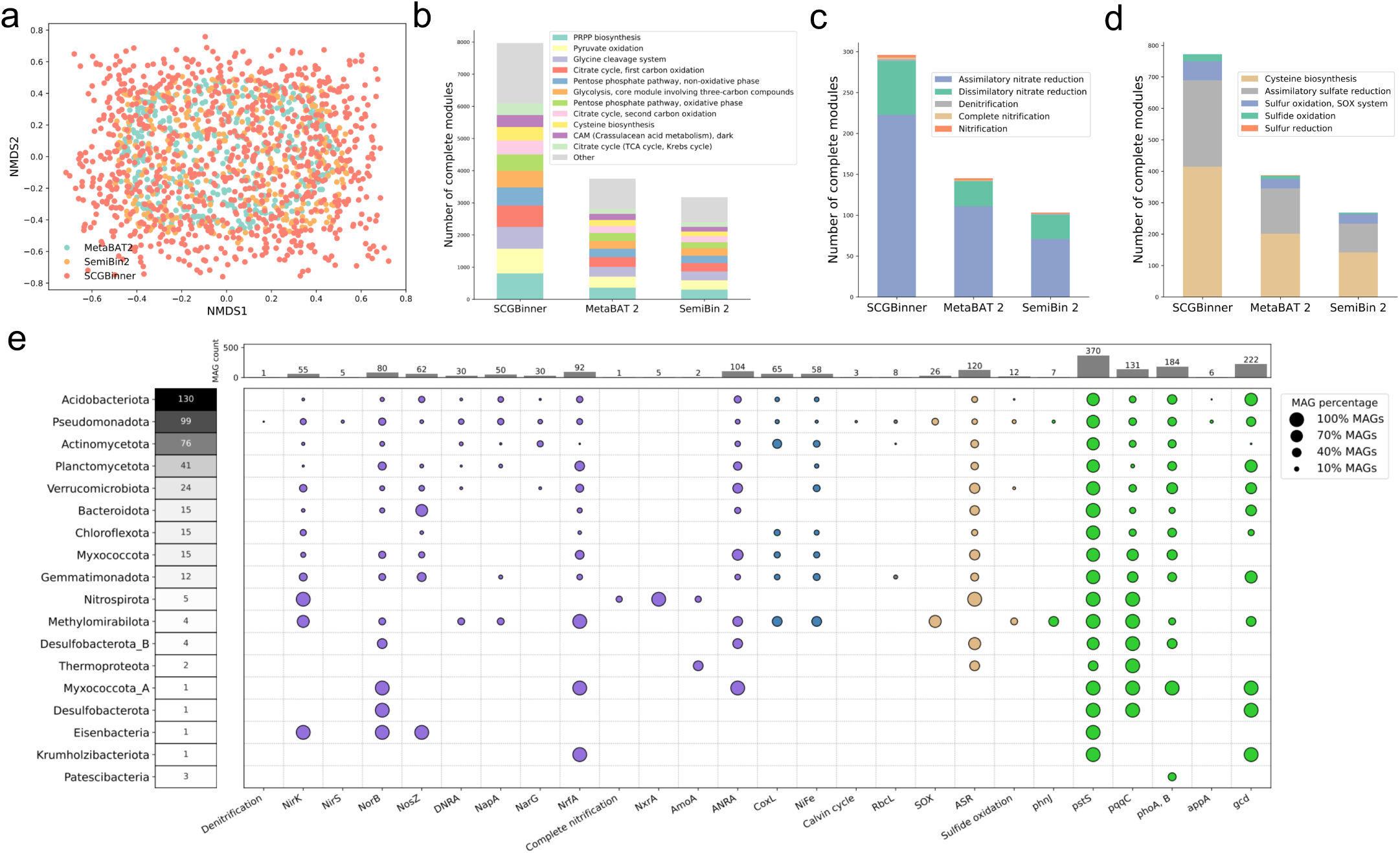
a, Non-metric multidimensional scaling (NMDS) based on binary Bray–Curtis dissimilarity of functional profiles (KEGG KO presence/absence) for species-level MAGs recovered by three binning tools. **b, c, d,** Number of complete carbon, nitrogen, and sulfur metabolic pathway modules across HQ species recovered by three binning tools. **e,** Key nitrogen, sulfur, phosphorus, and carbon cycling genes and pathways in unique novel HQ species recovered by SCGBinner. The heatmap shows the number of HQ species in each phylum, the scatter dots represent the percentage of HQ species in the corresponding phylum that contain each gene or pathway, and the bar plot indicates the number of HQ species containing each gene or pathway.

The functional potential of 449 novel HQ species (Fig. 4c) uniquely recovered by SCGBinner provided unprecedented insights into the functioning of the agricultural soil microbiome. They shed light on the predicted ecological roles of 65 uncharacterised families and 93 novel genera, increasing the resolution of microorganisms driving both the retention and loss of bioavailable nutrients essential for plant growth and productivity by between 40-100% relative to SemiBin2 and MetaBAT2 (1,042 species: 457 unique to SCGBinner, 467 overlapping species also recovered by SCGBinner, and 118 species recovered by SemiBin2 or MetaBAT2), including the cycling of trace and climate active gases (Fig. 6e).

These unique species highlight the integral role of the microbiome in mediating nitrogen bioavailability in soil, controlling the transformation of inorganic nitrogen, the exchange of *N*_2_, the release of climate active *N*_2_*O*, and the remineralisation of organic nitrogen for uptake by plants and other organisms (Fig. 6e). For instance, assimilatory nitrate reduction to ammonia (ANRA) stabilises nitrogen within soil through the incorporation of inorganic nitrogen into microbial biomass. A complete ANRA pathway was identified in 104 novel species unique to SCGBinner spanning 11 phyla, representing a 71% increase in the total number of species encoding this function (104 vs. 146 species, which include 2 unique known species and 109 overlapping novel species recovered by SCGBinner, and 35 novel species recovered by SemiBin2 or MetaBAT2). These 104 novel species belonged to 48 distinct families and 61 genera, of which 82% (50/61 genera) remain uncharacterised and are known only from environmental DNA sequences with placeholder taxonomic assignments.

Moreover, the unique species encoded several sulfur transformations controlling the availability of soil sulfur compounds for uptake by plants (Fig. 6e). We identified 26 MAGs encoding the SOX system for thiosulfate oxidation to sulfate, including 3 uncharacterised bacterial families of the SG8-39 and SG8-41 (Pseudomonadota) and CSP1-6 (Methylomirabilota), representing a 65% increase in the total number of MAGs encoding the SOX system across all binning tools. Sulfide oxidation, for the conversion of reduced sulfur compounds such as hydrogen sulfide back into elemental sulfur, was encoding by 12 MAGs including 3 uncharacterised families, UBA2999 (Acidobacteriota), CSP1-6 (Methylomirabilota), and JADJPG01 (Verrucomicrobiota) spanning 7 uncharacterised and one novel genera, indicative of an 86% increase in the recovery of microorganisms mediating this key process supporting sulfur regeneration.

Among the unique species, phosphorous cycling for both microbial growth and metabolism and regeneration within the soil environment was widespread (Fig. 6e). Alkaline phosphatase, which facilitates the dephosphorylation of organic compounds to release phosphate, encoded by *phoD*, was identified in 266 novel species unique to SCGBinner, equivalent to a 73% increase relative to the other binning tools. These MAGs spanned 13 phyla and were dominated by the UBA2999 and 2-12-FULL-66-21 families of the Acidobacteriota (50 and 20 MAGs respectively). Meanwhile, organic phosphorous mineralisation from plant-derived phytic acid, encoded by the *appA* gene, was highly specialised. Only 6 MAGs encoded *appA*, 5 from the Pseudomonadota and 1 of the Acidobacteriota. Nevertheless, this represented a 43% increase in the recovery of MAGs encoding this function. These MAGs belonged to 3 uncharacterised genera, Gp7-AA6 (Acidobacteriota), PALSA-894 (Pseudomonadota), and VBCG01 (Pseudomonadota), as well as the known genera *Reyranella* and *Sphingomicrobium* of the Pseudomonadota, indicative of crop rhizosphere microbiota.

### 2.8 Resource usage

The runtime and memory usage of COMEBin, LorBin, SemiBin2, and SCGBinner were evaluated on a machine equipped with two Intel(R) Xeon(R) Gold 6226R CPUs (2.90 GHz) and an NVIDIA GeForce RTX 3080 GPU. COMEBin, SemiBin2, and SCGBinner were executed with 64 threads and one GPU. LorBin was run using the default command because specifying the number of threads resulted in an error. Compared with SemiBin2, LorBin and COMEBin, SCGBinner exhibited shorter runtimes on both simulated marine and plant datasets, with memory usage remaining below 10 GB (Supplementary Table 12).

## 3 Discussion

In this study, we present SCGBinner, a metagenomic binning method designed for long-read sequencing data. SCGBinner uses SCG-guided contrastive learning to learn high-quality embeddings and then employs Leiden-based ensemble clustering to further enhance binning performance. SCGBinner outperformed state-of-the-art binning methods, including SemiBin2, COMEBin, LorBin, and MetaBAT2, on simulated long-read datasets generated using CAMISIM, with similar results observed on de novo-assembled simulated marine and plant datasets. However, all binners struggled to recover MAGs from the de novo-assembled strain-madness dataset, where extreme strain diversity posed substantial challenges for current long-read assemblers. Importantly, extensive benchmarking showed that SCGBinner consistently outperformed other binning methods on real-world datasets generated using both PacBio HiFi and ONT sequencing, with similar advantages observed on real-world short-read datasets. For real-world PacBio HiFi samples, two assemblers metaFlye and metaMDBG were used to assemble each sample, and seven binning methods were benchmarked on the assemblies. All binners recovered more “moderate-or-higher” quality, high-quality, and near-complete MAGs from metaMDBG assemblies than from metaFlye assemblies across soil, marine, sheep rumen, human gut, and AD sludge datasets. SCGBinner consistently achieved the best performance across all PacBio HiFi datasets for both metaMDBG and metaFlye assemblies. Notably, a significant positive correlation was observed between microbial diversity and the relative improvement in MAG recovery by SCGBinner compared to SemiBin2 across all PacBio HiFi samples, indicating that SCGBinner enhances MAG recovery from high-diversity microbial communities and improves our ability to understand the phylogenetic and functional diversity of notoriously difficult to assemble and bin microbial communities in soil and marine environments.

Single-copy genes (SCGs) are widely used in deep learning–based binning methods to facilitate MAG recovery. For instance, COMEBin, SemiBin, SemiBin2, and LorBin, utilize SCG information during clustering. However, existing state-of-the-art methods overlook the intrinsic advantages of long-read data when learning embeddings for the highly contiguous contigs. In this study, we observed that contigs assembled from long-read data contain a higher proportion of SCGs and more SCGs per contig. This enrichment allows SCGs to provide effective guidance for representation learning. SCGBinner improves current long-read binning methods by directly exploiting the advantages of long-read sequencing during contrastive learning, making it particularly suitable for real-world complex microbial communities.

Ensemble strategies are used in both bin-refinement tools and binning methods to integrate the strengths of multiple binning results. SemiBin2 uses SCGs to estimate F1-score, completeness, and contamination of bins across all clustering results, and then iteratively selects and updates bins under different contamination thresholds. SCGBinner improves upon the ensemble strategies of SemiBin2 by iteratively selecting and updating bins under stricter thresholds, and then outputs the bins in the optimal partition to ensure the recovery of potential MAGs that lack the SCGs used in this study (e.g., eukaryotic MAGs). Moreover, SCGBinner updates only the affected bins in each iteration to improve efficiency. SCGs serve as a guiding factor in representation learning and ensemble clustering, jointly contributing to the high performance of SCGBinner on complex real-world microbial communities (Supplementary Note 1). Presently, SCGBinner only utilises bacterial and archaeal SCGs. In future work, we plan to incorporate SCGs from other domains, such as eukaryotes, to further enhance SCGBinner’s performance.

In summary, SCGBinner leverages the advantages of long-read metagenomic sequencing and outperforms other state-of-the-art binning methods on both simulated and real-world datasets. Notably, it demonstrates outstanding performance in high-diversity real-world environments. When combined with continually improving long-read sequencing and superior long-read assembly approaches, SCGBinner is expected to serve as a powerful tool for analyzing metagenomic data, especially for surveying complex microbial communities.

## 4 Methods

### 4.1 Data augmentation

In the contrastive learning framework of SCGBinner, we employ a data augmentation procedure to transform a single contig into multiple views and maximize the agreement between the embeddings of these augmented views for representation learning [23]. Here, we generate augmented views from each contig based on the assumption proposed in our previous study [14] that sequential fragments derived from a single contig originate from the same genome. To ensure that the augmented views are sufficiently long and distinct, we first divide each original view into three equal-length subsequences to generate three augmented views (Fig. 1c). Next, we divide the same original view into two equal-length subsequences to generate two more augmented views [21]. Augmented views shorter than 1,000 bp are replaced by randomly sampled fragments of at least 1,000 bp from their corresponding original views. In total, we generate six views for each contig: one original view and five augmented views.

### 4.2 Binning modes

Three binning modes were used to benchmark the performance of the binning methods: co-assembly, single-sample, and multi-sample binning (co-binning). Co-assembly binning facilitates the recovery of low-abundance microorganisms [45] by first coassembling sequencing data from all samples and then binning the assembled contigs using coverage information across all samples. Single-sample binning assembles and bins each sample individually. Multi-sample binning (co-binning) improves binning performance compared with single-sample binning [18, 46] by independently assembling sequencing data from each sample while utilizing coverage profiles across all samples for binning.

### 4.3 Construct TNF and coverage vectors

TNF and coverage vectors serve as the initial representation for each contig. A sliding window of length four was applied to generate a 136-dimensional TNF feature per contig, as described in ref. [14].

Co-assembly or multi-sample binning produces a 2*M* -dimensional coverage vector for each contig, defined as follows:

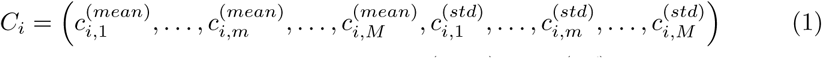

where *M* represents the number of samples, 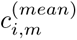 and 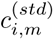 represent the mean and standard deviation of per-base coverage of the *i*-th contig by reads from the *m*-th sample, respectively. Each value in the coverage vectors is normalized across contigs after adding a small fraction.

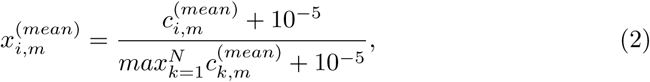

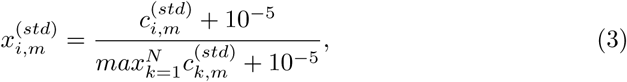

where *N* denotes the total number of contigs in a given sample. Define

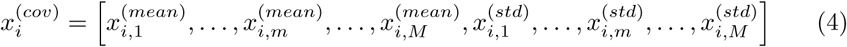

where 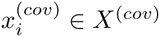 represents the coverage feature of *i*-th contig.

Single-sample binning produces a two-dimensional coverage vector for each contig, defined as follows:

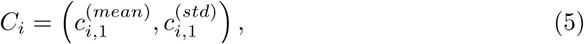

Each value in the coverage vectors is also normalized across contigs after adding a small fraction. The coverage feature of *i*-th contig is as follows:

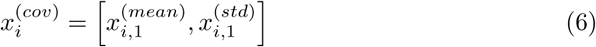

### 4.4 SCG guided contrastive learning

#### 4.4.1 Combined encoder

The combined encoder comprises two components: a coverage network and a combined network. Each component is implemented as a three-layer feed-forward neural network following the same configuration as COMEBin [14], with detailed parameter settings provided in Supplementary Table 13. Specifically, the coverage features (see Section “Construct TNF and coverage vectors”) are fed into the coverage network. The output of the coverage network is L2-normalized and concatenated with the TNF features to serve as input to the combined network. Finally, we applied L2 normalization to the output of the combined network to obtain embeddings for all contigs across all views.

#### 4.4.2 Average strategy for multiple views

Average strategy is used in this study for multi-view contrastive learning (Fig. 1d). Specifically, the embedding of each view and the average embedding of the other five views are treated as a positive pair to compute the loss function (see Section 4.4.4). This strategy facilitates faster convergence, as the augmented views may differ substantially [25].

#### 4.4.3 SCG batch

Contigs containing the same SCG harbor homologous coding regions with high sequence similarity, but they are expected to be assigned to different bins. Therefore, these contigs can be regarded as hard-negative pairs. To effectively exploit these hardnegative pairs, we build SCG batches for all SCGs in *G* = *{g*_1_*, … , g_k_, … , g_K_}*, where *K* = 107 denotes the total number of SCGs used in this study. For each SCG *g_k_*, we group the contigs containing *g_k_* into a training batch (SCG batch), where the batch size *N_g__k_* corresponds to the number of contigs carrying *g_k_*. For each view within a batch, the views of the other contigs in the same batch are treated as negative pairs to compute the contrastive loss. In addition, all contigs in a sample are split into batches of 1024 contigs (standard batches). Both SCG batches and standard batches are used in our training process (Supplementary Algorithm S1). The default value for the training epoch was set to 200.

#### 4.4.4 Objective function

The normalized temperature-scaled cross entropy loss (NT-Xent) [14, 23] is used to calculate the loss for each batch. Details of the training process can be found in Supplementary Algorithm S1. For a batch of data, the loss function is defined as:

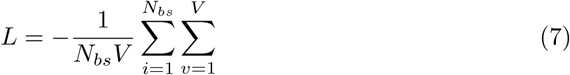

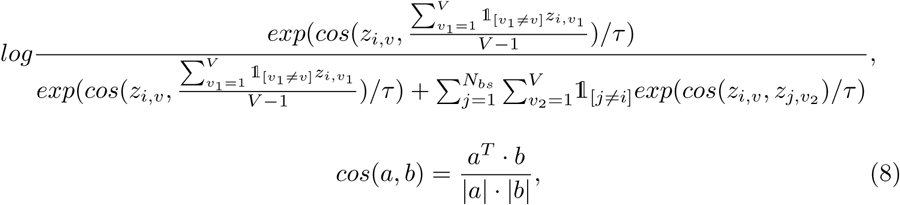

where *N_bs_*denotes the batch size, *V* (*V* = 6) represents the number of views for each contig, *z_i,v_* represents the embedding for *v*-th view of the *i*-th contig. *τ* is a temperature parameter, set to 0.07 for assemblies with an N50 greater than 10,000, and 0.15 otherwise [14]. For *z_i,v_*, the average embedding over the other five views of the *i*-th contig serves as its positive pair, while all views of the other contigs within the same batch are treated as negative pairs.

### 4.5 Ensemble clustering

After training the combined encoder using SCG-guided contrastive learning, the resulting contig embeddings were input into a Leiden-based ensemble clustering algorithm to generate the final non-redundant set of bins (see Supplementary Algorithm S2 and Fig. 1e). The Leiden algorithm [26] is a parameter-sensitive community detection method. Accordingly, SCGBinner performs Leiden in parallel across a range of parameter values, adopting the same parameter settings and initial membership configuration as used in COMEBin [14].

Then, we estimate the F1-score, completeness, and contamination for all bins larger than 200 kbp using 107 SCGs as the method described in SemiBin2 [21]. For each Leiden clustering result, we count the number of bins that meet three quality thresholds: completeness *>* 90%, *>* 70%, and *>* 50%, each with contamination *<* 5%. The clustering result with the largest total number of bins satisfying these criteria is designated as the optimal partition.

Based on different thresholds of completeness and contamination, the bin with the highest F1-score is selected. If multiple bins have the same F1-score, the one with the largest N50 is chosen. The contigs in the selected bin are then removed from all clustering results. Inspired by MAGScoT [27], only the F1 score, completeness, and contamination of bins affected by these removals are updated to improve computational efficiency. This selection and update process is repeated until no bins meet the minimum required thresholds for completeness and contamination. Finally, the remaining bins in the optimal partition are retained. Only bins larger than 200 kbp are considered.

### 4.6 Benchmarking datasets and preprocessing

The simulated marine (ten samples, 70 GB), plant (21 samples, 210 GB) and strain-madness (100 samples, 482 GB) long-read datasets were generated using CAMISIM (version 1.3) [28], based on 977, 894, and 408 reference genomes from the CAMI II challenge [24], respectively. For each sample, we used the genome abundances provided by CAMI II and simulated reads with a length of 12,000 bp and an error rate of 0.01% using the wgsim simulator supported by CAMISIM. CAMISIM then generated error-free contigs and determined the corresponding genome binning gold standards based on mappings of the simulated reads to the reference genomes. Consistent with the CAMI II challenge, plasmids and other circular elements were excluded from the evaluation. After filtering, the simulated marine, plant, and strain-madness datasets contained 775, 468, and 408 complete genomes, respectively.

Seven real-world metagenomic datasets covering soil, marine, sheep rumen, AD sludge, human gut, and activated sludge environments were used for benchmarking (Supplementary Table 14-19). The soil PacBio HiFi dataset contains one 250Gbp sample (the deep agricultural soil metagenome) [32]; the soil Oxford Nanopore dataset comprises the six samples with the largest read sizes (bp) selected from [33]; the marine dataset includes 30 PacBio HiFi and four Illumina samples [34]; the sheep rumen dataset contains one PacBio HiFi sample [35]; the AD sludge dataset includes three PacBio HiFi samples [15]; the human gut dataset contains four PacBio HiFi samples [15]; and the activated sludge dataset includes 23 Oxford Nanopore and 23 Illumina samples [36]. All datasets were used for benchmarking long-read single-sample binning, while the marine and activated sludge datasets were additionally used for long-read multi-sample binning, short-read single-sample binning, short-read multi-sample binning, hybrid single-sample binning, and hybrid multi-sample binning. The marine, human gut, and AD sludge datasets were also used for benchmarking long-read co-assembly binning.

The Illumina and Nanopore reads were quality-controlled and preprocessed using the same pipeline as described in ref. [18]. metaMDBG (version 1.2) [15] and metaFlye (version 2.9.2) [47] were used for single-sample assembly of PacBio HiFi reads. metaFlye (version 2.9.2) was used for single-sample assembly of Nanopore reads, the assembled contigs from Nanopore reads were further polished with Pilon [48]. For six Nanopore soil samples, the metagenomic assemblies and sequence data were obtained from ENA BioProject PRJEB58634. OPERA-MS (version 0.9.0) [16] was used for hybrid single-sample assembly. In hybrid single-sample assembly, each long-read sample and corresponding short-read sample were assembled together. For long-read samples obtained from ONT sequencing, the resulting hybrid assemblies were further polished using Pilon [48]. The resulting contigs longer than 1 kb were retained for binning. PacBio HiFi long reads and Illumina short reads were mapped to their corresponding contigs using minimap2 (version 2.24-r1171, -x map-hifi) and Bowtie2 (version 2.5.1), respectively. In hybrid data binning, when the hybrid assembly was generated from Illumina and Nanopore samples, only the short-read alignments were used as input for the binning process.

### 4.7 Compared methods and evaluation metrics

Several state-of-the-art binning methods were compared to our approach, including MetaBAT2 [8] (version 2.15), MetaBinner [11] (version 1.4.4), MetaDecoder [10] (version 1.0.16), SemiBin2 [21] (version 1.5.1), COMEBin [14] (version 1.0.3), and LorBin [22] (version 0.1.0).

We used CheckM 2 [49] (version 1.0.2) to assess the contamination and completeness of MAGs recovered from real-world datasets. For simulated datasets, we used AMBER (version 2.0.3) [29] to calculate the accuracy (bp), F1 score for sample (bp), and the number of recovered MAGs under different thresholds of contamination and completeness. We then define “moderate or higher” quality MAGs with contamination *<*10% and completeness *>*50%, high-quality MAGs with contamination *<*10% and completeness *>*70%, and near-complete MAGs with contamination *<*5% and completeness *>*90%. Pseudo-F statistics from permutational analysis of variance (PERMANOVA) were used to assess cluster separation performance, calculated using the skbio package in python.

### 4.8 Processing and analysis of MAGs

Nonpareil [37] (version 3.5.5) was used to evaluate the diversity and average coverage of PacBio HiFi samples. dRep [50] (version 3.4.3) was employed to cluster MAGs at the species level using the options -nc 0.6 -sa 0.95. GTDB-Tk [51] (version 2.4.0) with GTDB release r220 was employed to annotate species-level MAGs. IQ-TREE [52] (version 2.3.5) with the option -m LG+R4 was used to construct phylogenetic trees, which were subsequently visualized in iTOL v7 [53].

RGI (version 6.0.2) with default parameters was used to predict ARGs within MAGs, based on the Comprehensive Antibiotic Research Database [54] (CARD, version 3.2.7). Only ARGs present on contigs longer than 10 kb were considered [43]. MAGs containing more than one ARG type were classified as potential multi-resistance ARG hosts [44]. Potential BGCs in MAGs were annotated using antiSMASH [55] (v6.1.1), and their novelty was assessed with BiG-SLiCE [41] (v1.1.1) against 1,225,071 BGCs from the BiG-FAM database [40] at thresholds of 900 (default), 1200, and 1500. Higher thresholds indicate more stringent criteria, and BGCs with distances exceeding the threshold were considered potentially novel.

MAG functions were annotated using eggNOG-mapper (v2.1.13) [56] with the eggNOG database (v5.0.2) [57]. The completeness of metabolic modules was calculated using the kegg-pathways-completeness tool (https://github.com/EBI-Metagenomics/kegg-pathways-completeness-tool). Moreover, the predicted proteins from MAGs were mapped against a customized collection of 51 metabolic marker protein databases [58] using DIAMOND (v2.1.15) [59]. These marker proteins cover essential pathways for energy conservation from organic and inorganic compounds, aerobic and anaerobic respiration, nitrogen fixation, carbon fixation, and phototrophy. Gene hits were first filtered with at least 80% subject coverage or 80% query coverage and were further filtered using the same percentage identity thresholds as ref. [60].

### 4.9 Data availability

Sequence data from marine [34], sheep rumen [35], AD sludge [15], human gut [15], and activated sludge [36] samples are publicly available via the NCBI Sequence Read Archive. Sequence data from the deep agricultural soil metagenome are available via the ENA BioProject PRJEB88618 [32]. Sequence data and metagenomic assemblies of six Oxford Nanopore soil samples are publicly available from ENA BioProject PRJEB58634 [33]. The accession numbers and sample description are listed in Supplementary Table 14.

### 4.10 Code availability

The source code is available on GitHub (https://github.com/htaohan/SCGBinner) under the MIT License and is archived on Zenodo (https://doi.org/10.5281/zenodo.21998270).

## 5 Acknowledgements

This work has been supported by the National Natural Science Foundation of China (Grant Nos. U24A20257 and 62272105), ZJ Lab, Shanghai Center for Brain Science and Brain-Inspired Intelligence Technology, and 111 Project (Grant No. B18015).

## 6 Author contributions

S.Z. conceived the project. S.Z. and Z.W. supervised the project. S.Z., Z.W., and H.H. designed the study and the methodological framework. H.H. implemented the methods. S.Z., Z.W., and H.H. analyzed the benchmarking results. H.H., L.F.M., and S.R. analyzed the recovered genomes. L.F.M. analyzed the functional potential of MAGs. H.H. drafted the manuscript. S.Z., Z.W., H.H., L.F.M., C.Q., and G.D.B. revised the manuscript. C.Q. and S.R. provided the deep agricultural soil sample. All authors agree to the content of the final paper.

