## Supplementary file for "Advancing long-read metagenomic binning via single-copy-gene guided contrastive learning"

### Supplementary Note 1

The contribution of SCG-guided contrastive learning and ensemble clustering to binning performance was evaluated on the high-diversity real-world marine dataset (30 HiFi samples, Supplementary Fig. 8). The results showed that both SCG-guided contrastive learning and ensemble clustering countribute to the recovery of MAGs. Specifically, SCG-guided contrastive learning recovered an additional 323 and 74 near-complete MAGs based on contigs assembled with metaMDBG and metaFlye, respectively (SCGBinner-Ens compared to SCGBinner-Ens-SCG). Ensemble clustering further recovered 158 and 51 additional near-complete MAGs based on contigs assembled with metaMDBG and metaFlye, respectively (SCGBinner compared to SCGBinner-Ens). Notably, SCGBinner without ensemble clustering (SCGBinner-Ens) still recovered more high-quality and near-complete MAGs than other state-of-the-art binners, regardless of whether the contigs were assembled by metaMDBG or metaFlye.

### Supplementary Note 2

In addition to the simulated gold-standard assemblies generated by CAMISIM [1], we also generated simulated datasets by de novo assembling long reads produced with Badread [2]. Specifically, PacBio HiFi reads for the simulated marine (10 samples, 115 GB), plant (21 samples, 207 GB), and strain-madness (100 samples, 612 GB) datasets were generated using Badread (version 0.4.2). The simulated marine and plant datasets were separately co-assembled using metaMDBG (version 1.2) [3]. The simulated strain-madness dataset was co-assembled using metaMDBG (version 1.4), as metaMDBG version 1.2 failed to complete the assembly after one month of execution using 64 threads and 490 GB of memory. The generated contigs were then mapped back to the reference genomes, and the primary alignment with the largest number of matching bases was assigned as the ground truth genome for each contig. For MAG quality assessment, completeness was calculated as the fraction of the reference genome covered by the MAG, where the reference genome with the highest coverage breadth was considered the true source genome of the MAG. Contamination was calculated as the sum of (i) bases from contigs incorrectly assigned to other genomes and (ii) unmapped bases from correctly assigned contigs, divided by the bin size.

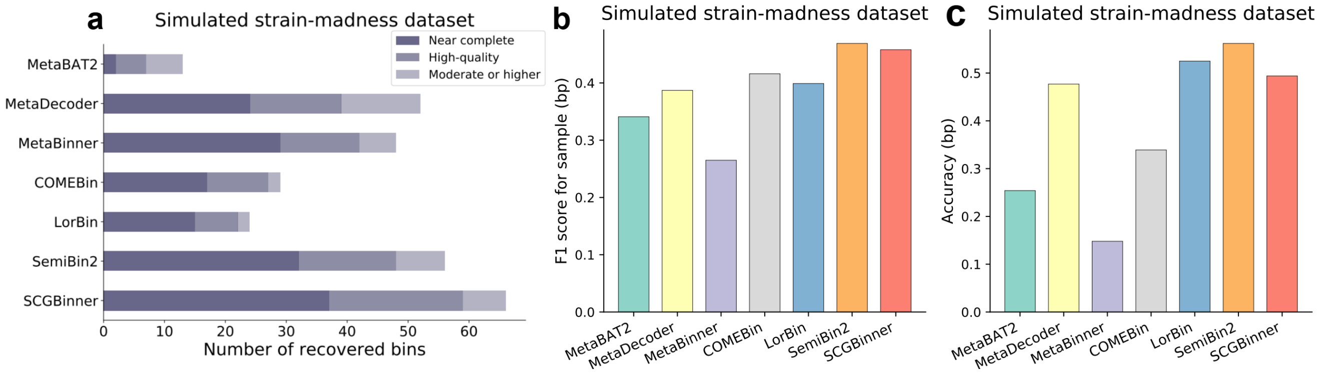


**Supplementary Figure 1** Comparison of binning performance on the simulated long-read strain-madness dataset generated using CAMISIM and containing 408 complete genomes. **a,** The number of recovered near complete, high-quality, and "moderate or higher" quality MAGs. **b,** F1 score for sample of different binning methods. **c,** Accuracy (bp) of different binning methods.

**Supplementary Table 1** Average percentage of contigs containing single-copy genes (SCGs), along with the overall mean of sample-wise average SCGs per contig (calculated among SCG-containing contigs). Each short-read sequencing sample was assembled individually using MEGAHIT. Each PacBio HiFi long-read sequencing sample was individually assembled using metaFlye and metaMDBG. Each Oxford Nanopore long-read sequencing sample was individually assembled using metaFlye.

| | Marine long-read dataset (metaFlye, 30 samples) | marine short-read dataset (4 samples) |
|---|---|---|
| Average percentage of contigs containing SCGs | 27.42% (±4.79) | 3.38% (±0.19) |
| Overall mean of sample-wise average SCGs per contig | 2.87 (±0.25) | 1.63 (±0.01) |
| | Marine long-read dataset (metaMDBG, 30 samples) | marine short-read dataset (4 samples) |
| Average percentage of contigs containing SCGs | 17.90% (±2.57) | 3.38% (±0.19) |
| Overall mean of sample-wise average SCGs per contig | 4.22 (±0.25) | 1.63 (±0.01) |
| | Activated sludge long-read dataset (metaFlye, 23 samples) | Activated sludge short-read dataset (23 samples) |
| Average percentage of contigs containing SCGs | 11.59% (±1.81) | 2.47% (±0.18) |
| Overall mean of sample-wise average SCGs per contig | 3.98 (±0.53) | 1.56 (±0.06) |

**Supplementary Table 2** Overview of the long-read simulated datasets generated by CAMISIM.

| | Plant | Marine | Strain-madness |
|---|---|---|---|
| Total length | 1,454,902,377 | 2,360,946,621 | 1,362,286,956 |
| # contigs | 37,470 | 32,677 | 15,633 |
| N50 | 72,755 | 206,083 | 150,534 |

**Supplementary Table 3** Overview of the long-read simulated datasets assembled using metaMDBG.

| | Plant | Marine | Strain-madness |
|---|---|---|---|
| Total length | 1,883,351,066 | 2,288,209,472 | 297,810,225 |
| # contigs | 23,116 | 11,298 | 3,429 |
| N50 | 195,574 | 1,529,430 | 217,531 |

**Supplementary Table 4** Permutational analysis of variance (PERMANOVA) results of contig embeddings from 15 randomly selected genomes (>100 contigs) in the simulated plant dataset.

| Binning method | PERMANOVA pseudo-F statistics | PERMANOVA $R^2$ value |
|---|---|---|
| SCGBinner | 586.99 | 0.72 |
| LorBin | 372.23 | 0.62 |
| COMEBin | 166.96 | 0.42 |

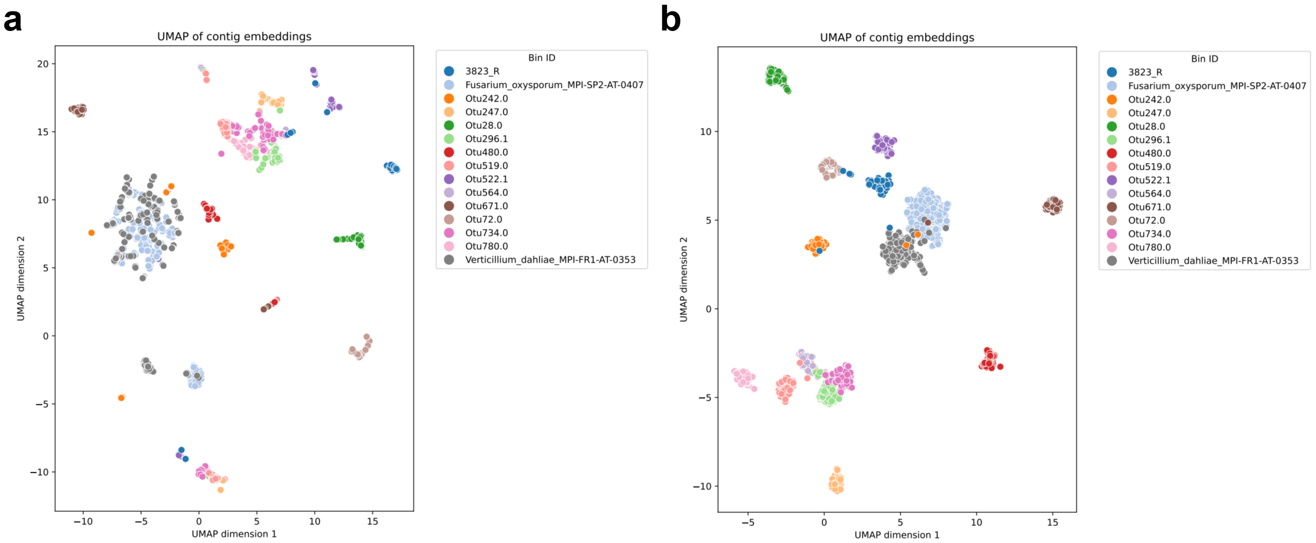


**Supplementary Figure 2** UMAP visualization of contig embedding on the simulated plant dataset. **a,** contig embeddings generated by LorBin. **b,** contig embeddings generated by COME-Bin.

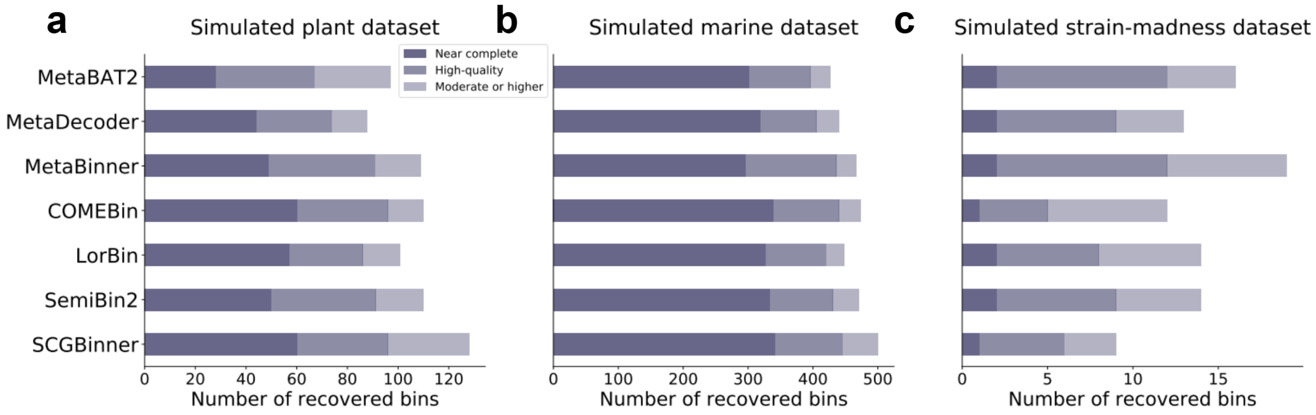


**Supplementary Figure 3** Binning performance on simulated marine, plant, and strain-madness datasets de novo assembled from long reads generated with Badread. **a,** Number of recovered near-complete, high-quality, and "moderate or higher" quality MAGs from the plant dataset. **b,** Number of recovered near-complete, high-quality, and "moderate or higher" quality MAGs from the marine dataset. **c,** Number of recovered near-complete, high-quality, and "moderate or higher" quality MAGs from the strain-madness dataset.

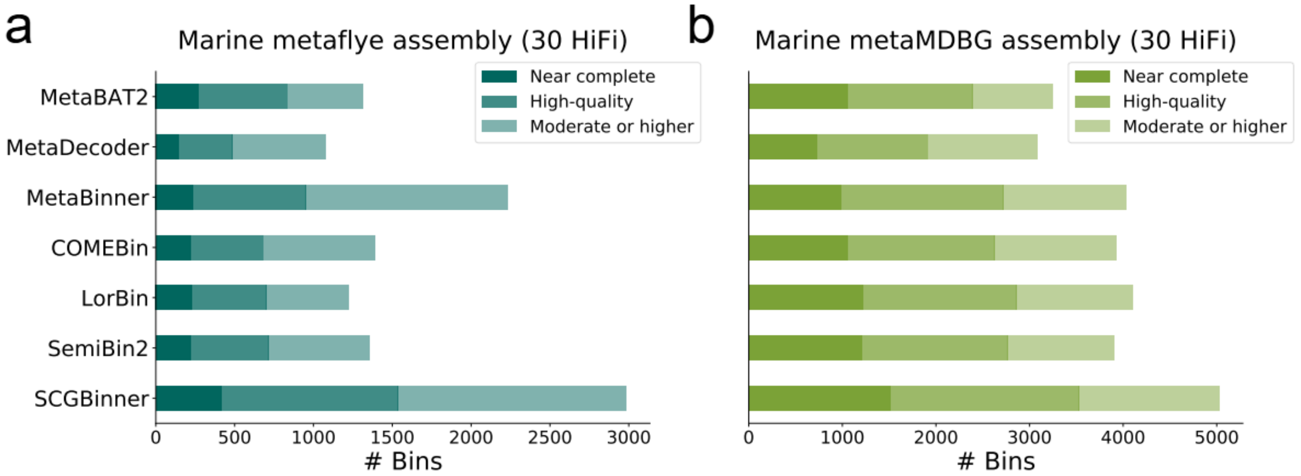


**Supplementary Figure 4** Comparison of binning methods on the marine dataset using multi-sample binning. **a,** The number of near-complete, high-quality, and "moderate or higher" quality MAGs recoverd from contigs assembled by metaflye. **b,** The number of near-complete, high-quality, and "moderate or higher" quality MAGs recoverd from contigs assembled by metaMDBG.

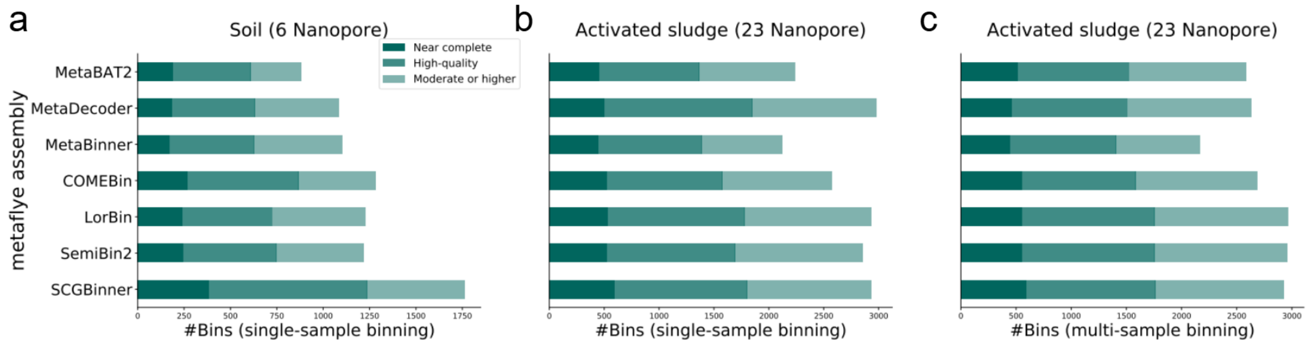


**Supplementary Figure 5** Comparison of binning methods on two Nanopore datasets. **a,** The number of near-complete, high-quality, and "moderate or higher" quality MAGs recovered from soil samples using single-sample binning. **b,** The number of near-complete, high-quality, and "moderate or higher" quality MAGs recovered from activated sludge samples using single-sample binning. **c,** The number of near-complete, high-quality, and "moderate or higher" quality MAGs recovered from activated sludge samples using multi-sample binning.

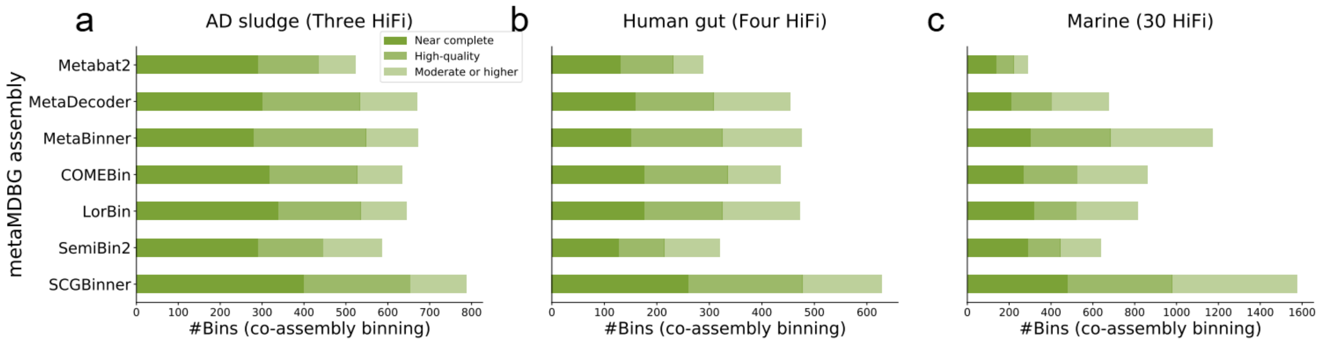


**Supplementary Figure 6** Comparison of binning methods on three real-world datasets using co-assembly binning with metaMDBG assembly. **a,** The number of near-complete, high-quality, and "moderate or higher" quality MAGs recovered from AD sludge samples using co-assembly binning. **b,** The number of near-complete, high-quality, and "moderate or higher" quality MAGs recovered from human gut samples using co-assembly binning. **c,** The number of near-complete, high-quality, and "moderate or higher" quality MAGs recovered from marine samples using co-assembly binning.

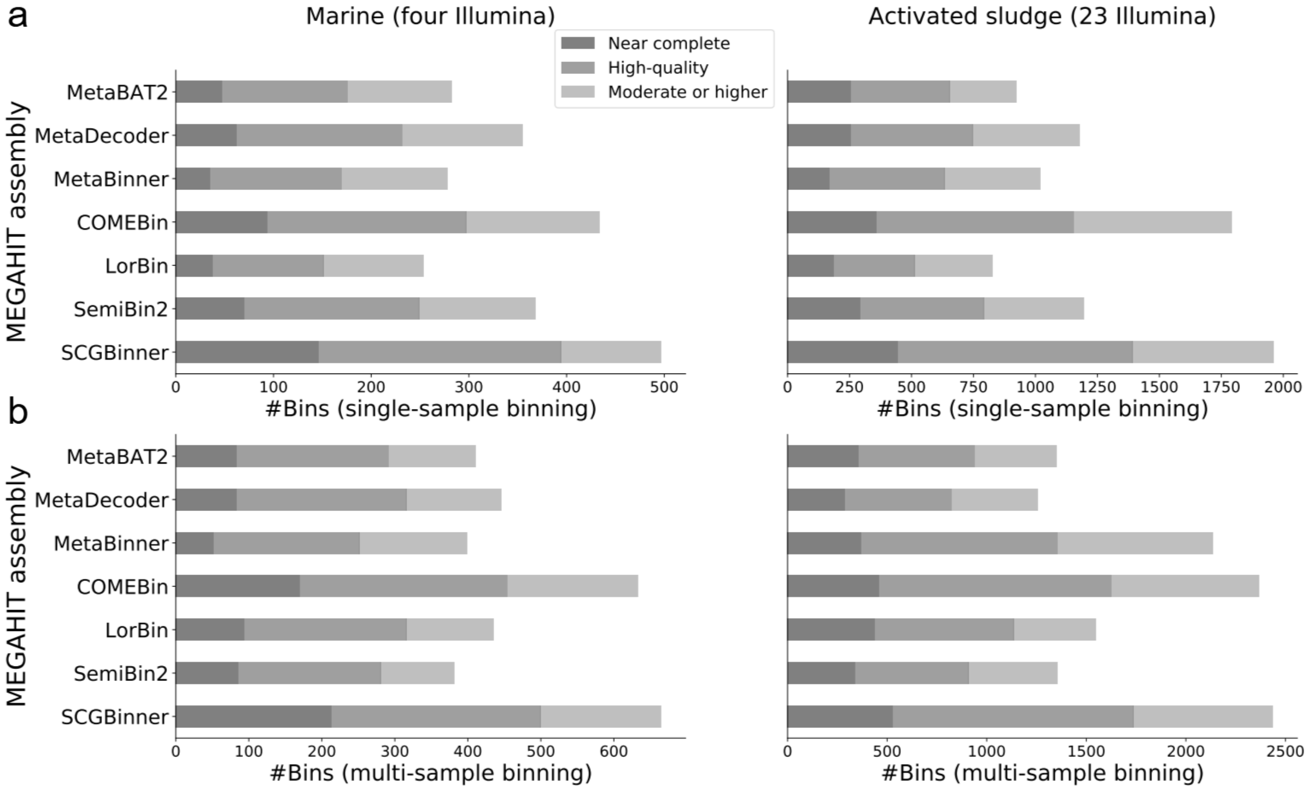


**Supplementary Figure 7** Comparison of binning methods on two real-world short-read datasets. **a,** The number of MAGs of near-complete, high-quality, and moderate-or-higher quality recovered from the marine and activated sludge datasets using single-sample binning. **b,** The number of MAGs of near-complete, high-quality, and moderate-or-higher quality recovered from the marine and activated sludge datasets using multi-sample binning.

---

**Algorithm S1** The single-copy-gene guided contrastive learning trainning process of SCGBinner

---

**Input:** $N_{sta}$: standard batch size. $V$: the number of views. $f(\cdot)$: combined encoder. $x_{i,v}^{com}, x_{i,v}^{cov}$ : TNF and coverage features for the $v$-th view of the $i$-th contig, respectively. $G$: all single-copy genes (SCGs) used to build SCG batches. $N_{g_k}$ : SCG batch size for $g_k$.

**Output:** combined encoder $f(\cdot)$

**for** each stadard batch $\left\{\left\{x_{i,v}^{(com)}, x_{i,v}^{(cov)}\right\}_{v=1}^{V}\right\}_{i=1}^{N_{sta}}$ **do**

$z_{i,v} = f\left(x_{i,v}^{(com)}, x_{i,v}^{(cov)}\right)$

$L = -\frac{1}{N_{sta}V}\sum_{i=1}^{N_{sta}}\sum_{v=1}^{V} log\frac{exp(cos(z_{i,v},\frac{\sum_{v_1=1}^{V}\mathbb{1}_{[v_1\neq v]}z_{i,v_1}}{V-1})/\tau)}{exp(cos(z_{i,v},\frac{\sum_{v_1=1}^{V}\mathbb{1}_{[v_1\neq v]}z_{i,v_1}}{V-1})/\tau)+\sum_{j=1}^{N_{sta}}\sum_{v_2=1}^{V}\mathbb{1}_{[j\neq i]}exp(cos(z_{i,v}, z_{j,v_2})/\tau)}$

    update the combined encoder $f(\cdot)$ to minimize $L$

**end for**

**for** $g_k$ in $G = \{g_1, \ldots, g_k, \ldots, g_K\}$ **do**

    **if** $N_{g_k} >= 128$ **then**

        build the SCG batch $\left\{\left\{x_{i,v}^{(com)}, x_{i,v}^{(cov)}\right\}_{v=1}^{V}\right\}_{i=1}^{N_{g_k}}$ for $g_k$

        $z_{i,v} = f\left(x_{i,v}^{(com)}, x_{i,v}^{(cov)}\right)$

        $L = -\frac{1}{N_{g_k}V}\sum_{i=1}^{N_{g_k}}\sum_{v=1}^{V} log\frac{exp(cos(z_{i,v},\frac{\sum_{v_1=1}^{V}\mathbb{1}_{[v_1\neq v]}z_{i,v_1}}{V-1})/\tau)}{exp(cos(z_{i,v},\frac{\sum_{v_1=1}^{V}\mathbb{1}_{[v_1\neq v]}z_{i,v_1}}{V-1})/\tau)+\sum_{j=1}^{N_{g_k}}\sum_{v_2=1}^{V}\mathbb{1}_{[j\neq i]}exp(cos(z_{i,v}, z_{j,v_2})/\tau)}$

    **end if**

    update the combined encoder $f(\cdot)$ to minimize $L_{g_k}$

**end for**

**return** the combined encoder $f(\cdot)$

---

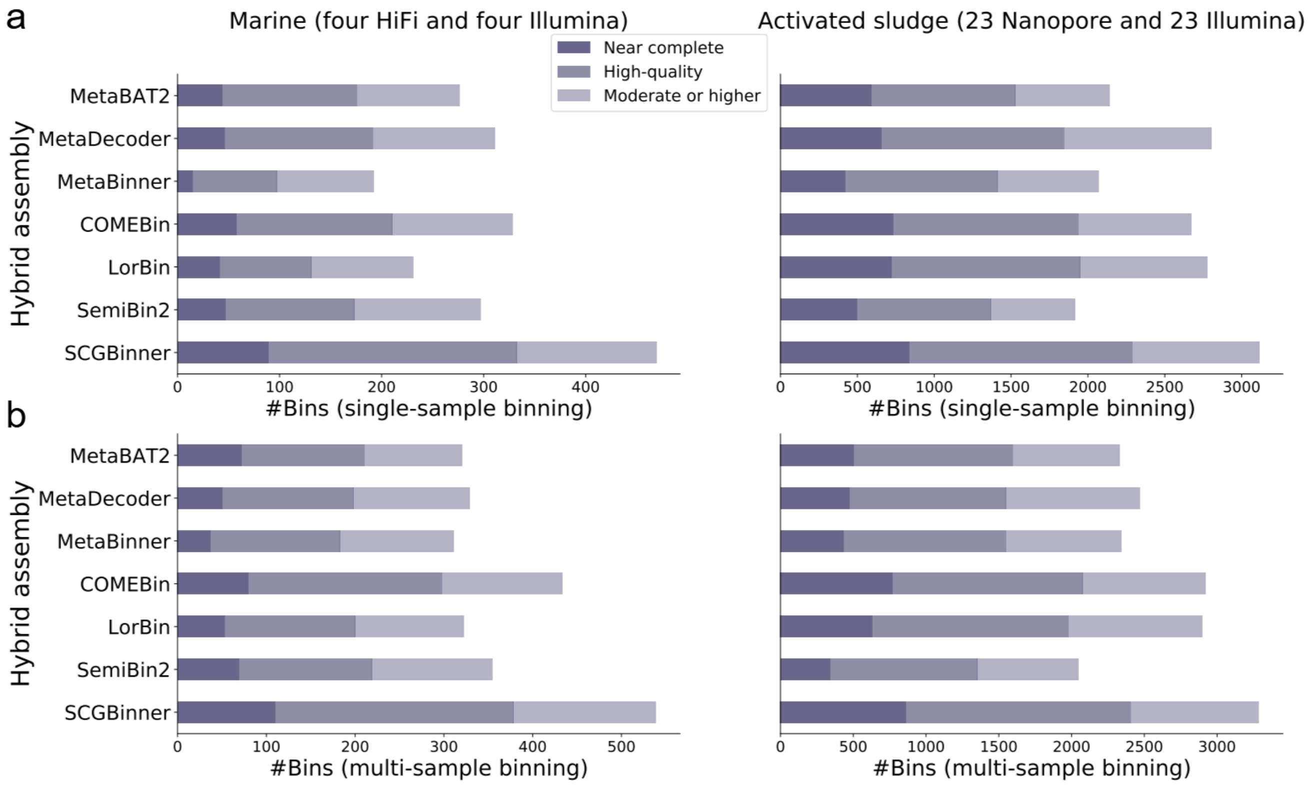


**Supplementary Figure 8** Comparison of binning methods on two real-world hybrid datasets. **a,** The number of MAGs of near-complete, high-quality, and moderate-or-higher quality recovered from the marine and activated sludge datasets using single-sample binning. **b,** The number of MAGs of near-complete, high-quality, and moderate-or-higher quality recovered from the marine and activated sludge datasets using multi-sample binning.

---

**Algorithm S2** Leiden-based ensemble clustering

---

**Input:** Contig embeddings $z_1, z_2, ..., z_n$; The Best_bin function returns the bin with the highest F1-score that meets the specified completeness and contamination thresholds. The Update function first removes contigs in the selected bin from $L_1$–$L_{120}$, then recalculates the F1-score, completeness, and contamination of the affected bins.

**Output:** Non-redundant bin set

```
Run Leiden with different parameter settings to obtain clustering results: L_1, L_2, ..., L_120.
Estimate the F1-score, completeness, and contamination for all bins larger than 200 kbp in L_1–L_120.
Obtain the optimal partition L_b
conta_threshold = 0.05
for comp_threshold= 0.9, 0.7, 0.5 do
    while bin is not None do
        bin=Best_bin(conta_threshold, comp_threshold)
        if bin is not None then
            output the returned bin;
            Update(bin, L_1, L_2, ..., L_120)
        end if
    end while
end for
Output the remaining bins larger than 200kbp in the optimal partition L_b
```

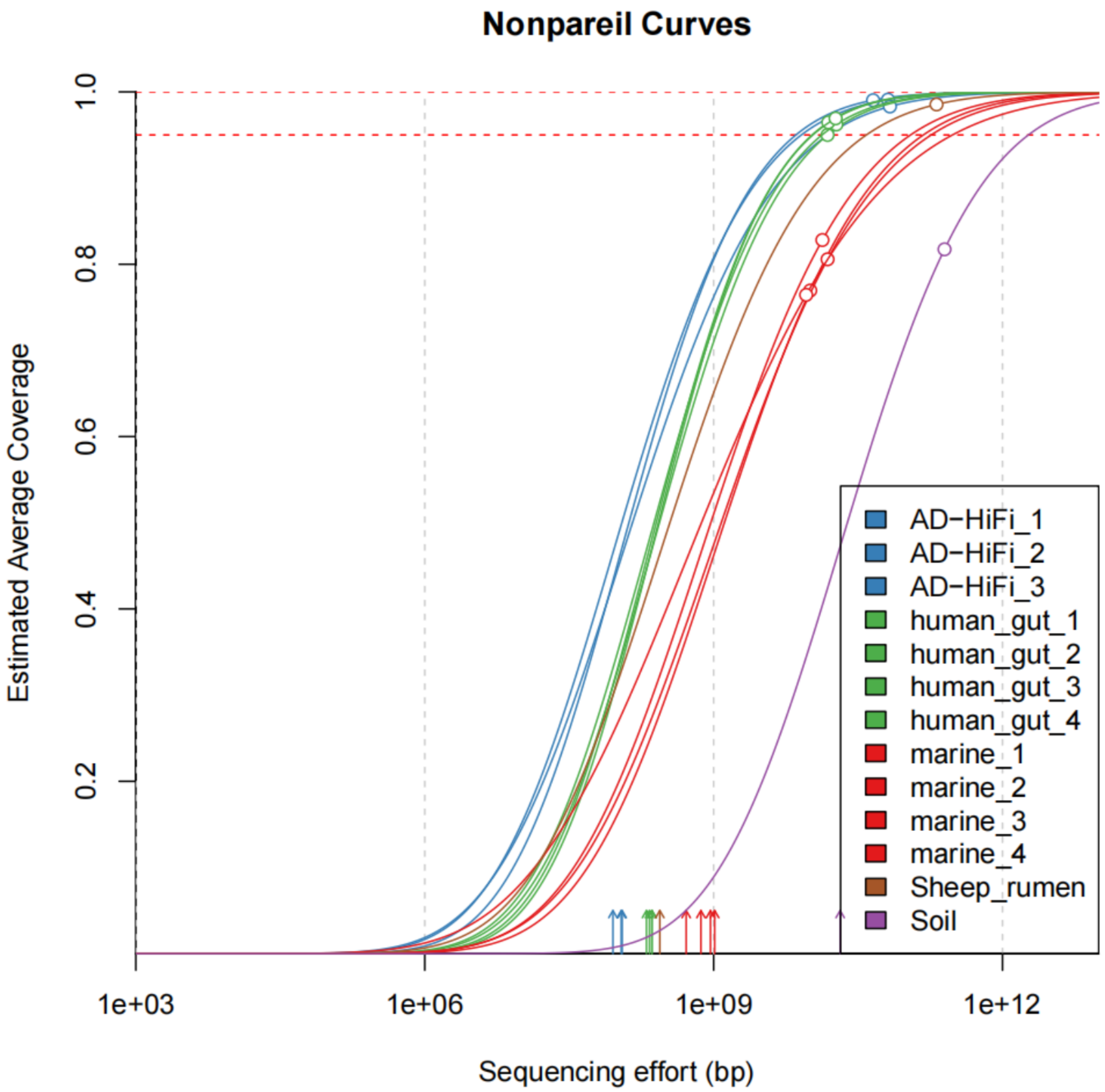


**Supplementary Figure 9** Nonpareil curves of samples in different enviroments.

**Supplementary Table 5** Number of NC MAGs containing the 23S, 16S, and 5S rRNA genes and at least 18 tRNAs recovered by different binning methods from real-world datasets assembled with metaFlye.

| | Soil sinlge-sample binning (one HiFi) | Marine single-sample binning (30 HiFi) | Marine multi-sample binning (30 HiFi) | Sheep rumen single-sample binning (one HiFi) | Human gut single-sample binning (four HiFi) | AD sludge single-sample binning (three HiFi) | Soil single-sample bin-ning (six Nanopore) | Activated sludge single-sample binning (23 Nanopore) | Activated sludge multi-sample binning (23 Nanopore) |
|---|---|---|---|---|---|---|---|---|---|
| MetaBAT2 | 10 | 76 | 256 | 107 | 80 | 80 | 112 | 354 | 402 |
| MetaDecoder | 2 | 87 | 129 | 194 | 110 | 131 | 118 | 405 | 377 |
| MetaBinner | 15 | 131 | 232 | 245 | 178 | 149 | 113 | 393 | 385 |
| COMEBin | 10 | 126 | 197 | 127 | 126 | 114 | 145 | 402 | 441 |
| SemiBin2 | 9 | 147 | 189 | 155 | 115 | 154 | 138 | 378 | 398 |
| LorBin | 12 | 127 | 199 | 152 | 107 | 142 | 188 | 392 | 403 |
| SCGBinner | 21 | 263 | 383 | 268 | 207 | 226 | 197 | 458 | 456 |

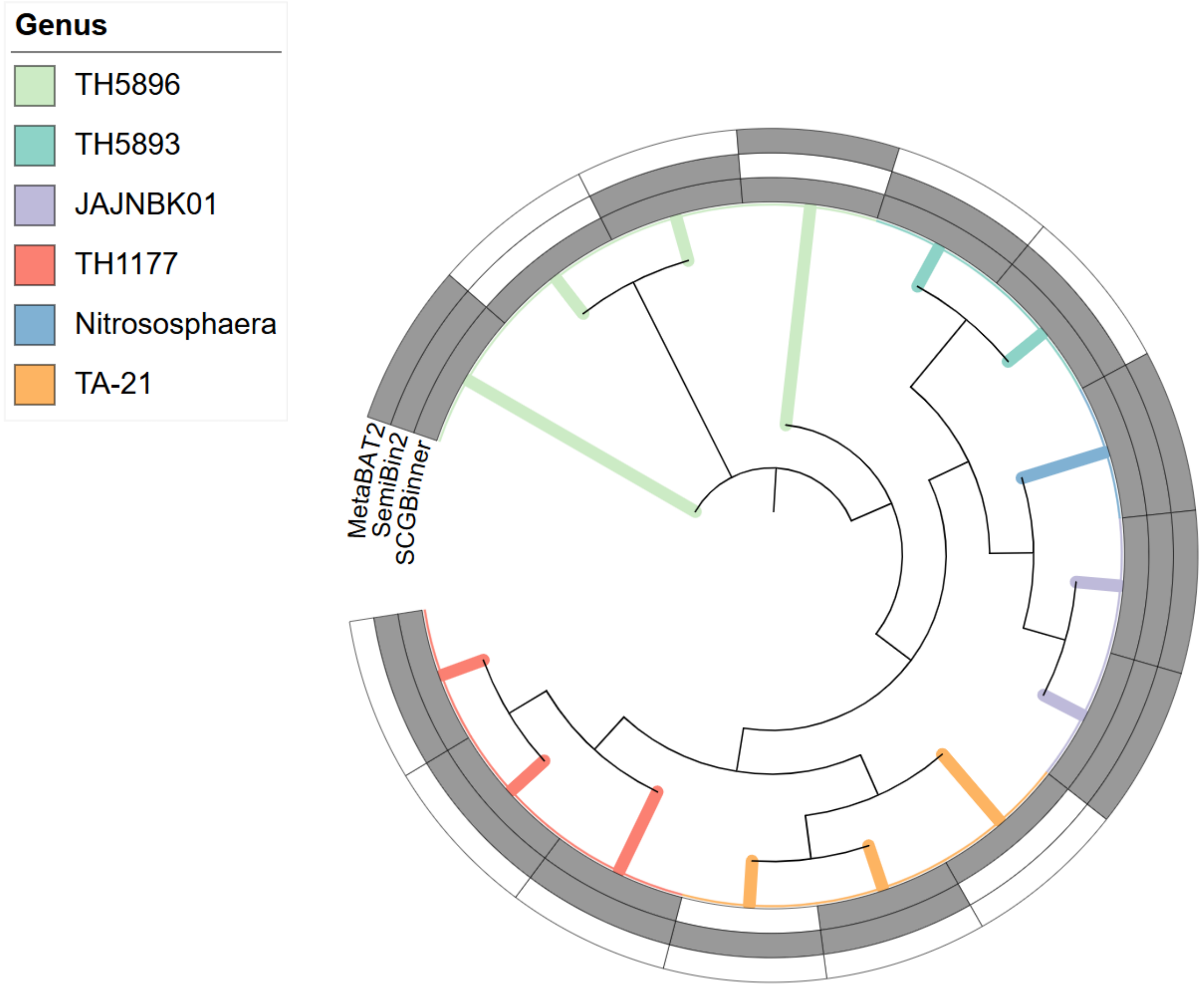


**Supplementary Figure 10** Phylogenetic tree of species-level HQ MAGs from a deep agricultural soil metagenome using GTDB-Tk archaeal marker genes.

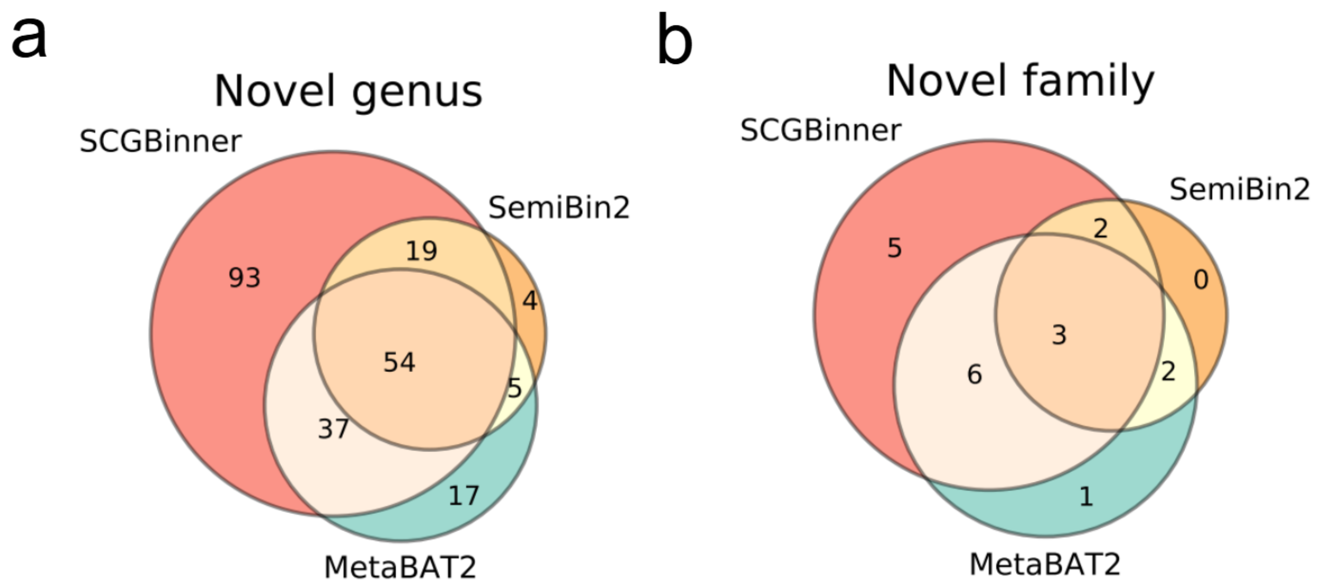


**Supplementary Figure 11** GTDB annotation of HQ species recovered from a deep agricultural soil metagenome. **a,** Number of novel genus per binner. **b,** Number of novel family per binner.

**Supplementary Table 6** Number of NC MAGs containing the 23S, 16S, and 5S rRNA genes and at least 18 tRNAs recovered by different binning methods from real-world datasets assembled with metaMDBG. "nan" denotes that the corresponding binners failed to produce binning results. COMEBin failed due to memory-related errors when executed with 16 threads on a machine with 503 GB of RAM, whereas LorBin was unable to complete the binning process on the same machine after one month of execution.

| | Soil sinlge-sample binning (one HiFi) | Marine single-sample binning (30 HiFi) | Marine multi-sample binning (30 HiFi) | Marine co-assembly binning (30 HiFi) | Sheep rumen single-sample binning (one HiFi) | Human gut single-sample binning (four HiFi) | Human gut co-assembly binning (four HiFi) | AD sludge single-sample binning (three HiFi) | AD sludge co-assembly binning (three HiFi) |
|---|---|---|---|---|---|---|---|---|---|
| MetaBAT2 | 92 | 480 | 857 | 110 | 148 | 118 | 63 | 239 | 165 |
| MetaDecoder | 73 | 590 | 561 | 108 | 252 | 190 | 88 | 308 | 196 |
| MetaBinner | 122 | 780 | 896 | 218 | 393 | 244 | 109 | 381 | 244 |
| COMEBin | nan | 522 | 800 | 52 | 178 | 152 | 83 | 230 | 197 |
| SemiBin2 | 74 | 565 | 756 | 75 | 192 | 173 | 30 | 276 | 112 |
| LorBin | nan | 560 | 780 | 109 | 200 | 164 | 70 | 264 | 163 |
| SCGBinner | 120 | 953 | 1051 | 256 | 312 | 236 | 141 | 342 | 223 |

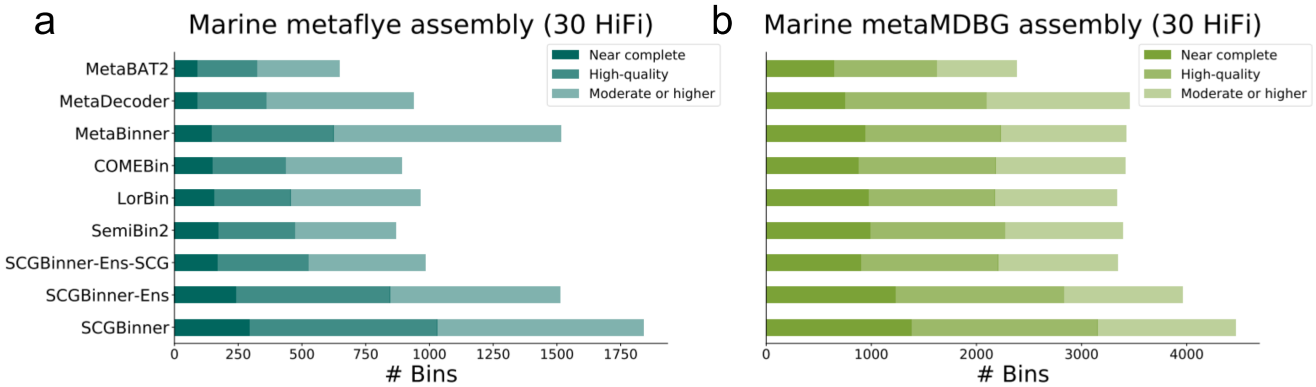


**Supplementary Figure 12** Effect of ensemble clustering and SCG-guided contrastive learning on the marine dataset using single-sample binning. **a,** The number of near-complete, high-quality, and "moderate or higher" quality MAGs recoverd from contigs assembled by metaflye. **b,** The number of near-complete, high-quality, and "moderate or higher" quality MAGs recoverd from contigs assembled by metaMDBG. SCGBinner-Ens refers to SCGBinner without ensemble clustering (the final binning result corresponds to the optimal partition). SCGBinner-Ens-SCG refers to SCGBinner without ensemble clustering and without SCG-guided contrastive learning (no SCG batches were used during training).

**Supplementary Table 7** Number of novel BGCs in HQ species recovered from a deep agricultural soil metagenome across different BGC types.

| BGC type | SCGBinner | SemiBin2 | MetaBAT2 |
|---|---|---|---|
| terpene | 748 | 243 | 388 |
| NRPS-like | 580 | 249 | 264 |
| T3PKS | 252 | 83 | 147 |
| RRE-containing | 205 | 94 | 100 |
| NRPS | 201 | 69 | 93 |
| T1PKS | 154 | 47 | 93 |
| betalactone | 145 | 72 | 78 |
| arylpolyene | 130 | 45 | 63 |
| RiPP-like | 94 | 41 | 54 |
| lassopeptide | 86 | 46 | 33 |
| ranthipeptide | 75 | 26 | 28 |
| redox-cofactor | 75 | 25 | 41 |
| phosphonate | 48 | 19 | 26 |
| thioamitides | 44 | 20 | 18 |
| other | 42 | 10 | 19 |
| ladderane | 42 | 9 | 20 |
| hglE-KS | 38 | 11 | 22 |
| acyl_amino_acids | 38 | 17 | 17 |
| resorcinol | 33 | 8 | 19 |
| NAPAA | 32 | 7 | 17 |
| indole | 30 | 8 | 8 |
| thiopeptide | 23 | 12 | 8 |
| oligosaccharide | 16 | 3 | 8 |
| hserlactone | 16 | 6 | 10 |
| lanthipeptide-class-v | 15 | 8 | 6 |
| LAP | 14 | 5 | 10 |
| phenazine | 12 | 6 | 4 |
| lanthipeptide-class-i | 11 | 3 | 6 |
| proteusin | 10 | 6 | 8 |
| nucleoside | 7 | 3 | 6 |
| PKS-like | 7 | 1 | 2 |
| transAT-PKS | 6 | 1 | 1 |
| lanthipeptide-class-iv | 6 | 5 | 4 |
| ectoine | 5 | 2 | 4 |
| T2PKS | 5 | 0 | 1 |
| siderophore | 5 | 1 | 2 |
| lanthipeptide-class-ii | 4 | 0 | 3 |
| cyanobactin | 4 | 1 | 2 |
| butyrolactone | 3 | 0 | 1 |
| thioamide-NRP | 3 | 3 | 1 |
| microviridin | 2 | 1 | 1 |
| bottromycin | 2 | 2 | 1 |
| linaridin | 2 | 1 | 0 |
| NAGGN | 2 | 1 | 4 |
| lipolanthine | 1 | 0 | 0 |
| aminocoumarin | 1 | 0 | 0 |
| furan | 1 | 1 | 1 |
| tropodithietic-acid | 1 | 2 | 0 |
| lanthipeptide-class-iii | 1 | 0 | 1 |
| sactipeptide | 1 | 0 | 0 |

**Supplementary Table 8** Number of unique novel BGCs identified in HQ species recovered from a deep agricultural soil metagenome across phyla.

| Phylum | SCGBinner | SemiBin2 | MetaBAT2 |
|---|---|---|---|
| Acidobacteriota | 430 | 43 | 83 |
| Pseudomonadota | 318 | 48 | 60 |
| Actinomycetota | 273 | 14 | 60 |
| Planctomycetota | 166 | 3 | 31 |
| Verrucomicrobiota | 132 | 0 | 23 |
| Myxococcota | 82 | 1 | 7 |
| Bacteroidota | 47 | 3 | 17 |
| Methylomirabilota | 40 | 0 | 8 |
| Chloroflexota | 34 | 2 | 22 |
| Nitrospirota | 31 | 5 | 0 |
| Desulfobacterota_B | 30 | 0 | 6 |
| Gemmatimonadota | 26 | 3 | 2 |
| Desulfobacterota | 7 | 0 | 0 |
| Myxococcota_A | 5 | 0 | 5 |
| UBA10199 | 2 | 0 | 0 |
| Thermoproteota | 2 | 2 | 0 |
| Eisenbacteria | 1 | 0 | 0 |
| Krumholzibacteriota | 1 | 0 | 0 |
| Armatimonadota | 0 | 0 | 4 |

**Supplementary Table 9** Number of complete carbon metabolism pathway modules in HQ species from a deep agricultural soil metagenome.

| Carbon metabolism | SCGBinner | MetaBAT2 | SemiBin2 |
|---|---|---|---|
| PRPP biosynthesis | 812 | 362 | 307 |
| Pyruvate oxidation | 762 | 345 | 288 |
| Glycine cleavage system | 686 | 313 | 272 |
| Citrate cycle, first carbon oxidation | 663 | 299 | 266 |
| Pentose phosphate pathway, non-oxidative phase | 568 | 261 | 231 |
| Glycolysis, core module involving three-carbon compounds | 507 | 242 | 223 |
| Pentose phosphate pathway, oxidative phase | 500 | 255 | 197 |
| Citrate cycle, second carbon oxidation | 447 | 203 | 182 |
| Cysteine biosynthesis | 414 | 201 | 142 |
| CAM (Crassulacean acid metabolism), dark | 375 | 178 | 145 |
| Citrate cycle (TCA cycle, Krebs cycle) | 365 | 164 | 156 |
| Phosphate acetyltransferase-acetate kinase pathway | 350 | 175 | 145 |
| Pentose phosphate pathway (Pentose phosphate cycle) | 347 | 180 | 154 |
| Glycolysis (Embden-Meyerhof pathway) | 300 | 160 | 142 |
| Serine biosynthesis | 258 | 123 | 91 |
| Glyoxylate cycle | 210 | 95 | 91 |
| Propanoyl-CoA metabolism | 156 | 81 | 54 |
| CAM (Crassulacean acid metabolism), light | 145 | 79 | 47 |
| Entner-Doudoroff pathway | 58 | 26 | 24 |
| Formaldehyde assimilation, ribulose monophosphate pathway | 13 | 5 | 5 |
| Reductive pentose phosphate cycle (Calvin cycle) | 11 | 5 | 5 |
| Pentose phosphate pathway | 8 | 3 | 6 |
| Formaldehyde assimilation, serine pathway | 5 | 0 | 0 |
| Ethylmalonyl pathway | 4 | 3 | 2 |
| Methylcitrate cycle | 1 | 2 | 1 |
| Reductive acetyl-CoA pathway (Wood-Ljungdahl pathway) | 1 | 0 | 0 |

**Supplementary Table 10** Number of complete nitrogen metabolism pathway modules in HQ species from a deep agricultural soil metagenome.

| Carbon metabolism | SCGBinner | MetaBAT2 | SemiBin2 |
|---|---|---|---|
| Assimilatory nitrate reduction | 223 | 111 | 71 |
| Dissimilatory nitrate reduction | 66 | 31 | 30 |
| Denitrification | 2 | 0 | 1 |
| Complete nitrification | 2 | 1 | 0 |
| Nitrification | 3 | 2 | 1 |

**Supplementary Table 11** Number of complete sulfur metabolism pathway modules in HQ species from a deep agricultural soil metagenome.

| Carbon metabolism | SCGBinner | MetaBAT2 | SemiBin2 |
|---|---|---|---|
| Cysteine biosynthesis | 414 | 201 | 142 |
| Assimilatory sulfate reduction | 275 | 145 | 91 |
| Sulfur oxidation, SOX system | 61 | 30 | 31 |
| Sulfide oxidation | 22 | 10 | 4 |
| Sulfur reduction | 0 | 1 | 0 |

**Supplementary Table 12** Runtime and memory usage of metagenomic binning tools on marine and plant datasets simulated using CAMISIM.

| Dataset | Simulated marine | Simulated plant |
|---|---|---|
| | Time (minutes) | |
| COMEBin | 106 | 92 |
| LorBin | 166 | 82 |
| SemiBin 2 | 80 | 64 |
| SCGBinner | 73 | 46 |
| | Memory (GB) | |
| COMEBin | 5.07 | 6.47 |
| LorBin | 4.04 | 4.85 |
| SemiBin 2 | 25.61 | 26.67 |
| SCGBinner | 8.05 | 6.28 |
| # Contigs | 32677 | 37470 |
| # Samples used for coverage information | 10 | 21 |

**Supplementary Table 13** Configuration of the coverage and combined networks.

| | #hidden layer | #hidden units | Input size | Output size | Use batch normalization | Activation function |
|---|---|---|---|---|---|---|
| Coverage network | 3 | 2048 | #samples | 128 | ✓ | LeakyReLu |
| Combined network | 3 | 2048 | 264 | 128 | ✓ | LeakyReLu |

"#hidden layers" means the number of hidden layers. "#hidden units" means the number of hidden units. "#samples" means the number of samples used to calculate the coverage feature for each contig.

**Supplementary Table 14** Run accessions and sample description for benchmarking real-world datasets.

| Dataset | Run accessions |
|---|---|
| Marine (Illumina) | ERR9769375 ERR9769376 ERR9769377 ERR9769378 |
| Marine (HiFi) | ERR9769275 ERR9769276 ERR9769277 ERR9769278 ERR9769279 ERR9769280 ERR9769281 ERR9769282 ERR9769283 ERR9769284 ERR9769285 ERR9769286 ERR9769287 ERR9769288 ERR9769289 ERR9769290 ERR9769291 ERR9769292 ERR9769293 ERR9769294 ERR9769295 ERR9769296 ERR9769297 ERR9769298 ERR9769299 ERR9769300 ERR9769301 ERR9769302 ERR9769303 ERR9769304 |
| Activated sludge (Illumina) | SRR11673989 SRR11673992 SRR11673995 SRR11673999 SRR11674002 SRR11674005 SRR11674008 SRR11674012 SRR11674015 SRR11674018 SRR11674022 SRR11674025 SRR11674031 SRR11674035 SRR11674038 SRR11674041 SRR11674044 SRR11674048 SRR11674051 SRR11674054 SRR11673976 SRR11674009 SRR11674045 |
| Activated sludge (Nanopore) | SRR11673963 SRR11673964 SRR11673966 SRR11673967 SRR11673968 SRR11673969 SRR11673970 SRR11673971 SRR11673972 SRR11673973 SRR11673974 SRR11673975 SRR11673977 SRR11673978 SRR11673979 SRR11673980 SRR11673981 SRR11673982 SRR11673983 SRR11673984 SRR11673985 SRR11673986 SRR11673988 |
| AD-HiFi (HiFi) | ERR10905741 ERR10905742 ERR10905743 |
| Sheep rumen (HiFi) | SRR14289618 |
| Human gut (HiFi) | SRR15275213 SRR15275212 SRR15275211 SRR15275210 |
| Soil (Nanopore) | ERR11718635 ERR12040030 ERR12036784 ERR11718641 ERR11594400 ERR11593883 |
| Soil (HiFi) | ERR15289804 |
| **Dataset** | **Sample Description** |
| Marine | Surface seawater samples collected from the North Sea between March and May 2020 [4], including 30 HiFi samples collected at 30 time points and four Illumina samples collected at four time points. |
| Activated sludge | Fresh activated sludge samples collected from 23 Danish wastewater treatment plants during August and September of 2018 [5] (23 mNGS samples and 23 Nanopore samples). |
| AD-HiFi (HiFi) | Time-series of three samples extracted from anaerobic digester sludge (week t+1, t+20 and t+40) [3]. |
| Sheep rumen (HiFi) | Sheep rumen microbiome [6]. |
| Human gut (HiFi) | Four pools each pool comprising samples from four vegan or omnivore individuals [3]. |
| Soil (HiFi) | Agricultural field in Oxfordshire (UK) [7]. |
| Soil (Nanopore) | The six Nanopore long-read soil metagenomic sequencing samples with the largest read sizes (bp) were selected from [8]. |

**Supplementary Table 15** Overview of metaFlye-based PacBio HiFi single-sample assemblies used for benchmarking.

| | Soil | Marine | Sheep rumen | Human gut | anaerobic digester sludge |
|---|---|---|---|---|---|
| # contigs (mean) | 590,850 | 40,761 | 51,521 | 14,514 | 42,245 |
| # contigs (max) | 590,850 | 68,585 | 51,521 | 15,273 | 47,161 |
| # contigs (min) | 590,850 | 16,061 | 51,521 | 13,265 | 37,307 |
| N50 (mean) | 35,170 | 27,885 | 212,979 | 143,000 | 56,468 |
| N50 (max) | 35,170 | 38,185 | 212,979 | 172,799 | 60,268 |
| N50 (min) | 35,170 | 17,068 | 212,979 | 117,215 | 52,511 |
| Total length (mean) | 17,196,825,394 | 890,945,443 | 4,322,828,271 | 1,031,057,120 | 1,532,761,312 |
| Total length (max) | 17,196,825,394 | 1,463,531,463 | 4,322,828,271 | 1,117,921,338 | 1,762,574,725 |
| Total length (min) | 17,196,825,394 | 340,495,661 | 4,322,828,271 | 923,733,920 | 1,320,051,275 |

**Supplementary Table 16** Overview of metaMDBG-based PacBio HiFi single-sample assemblies used for benchmarking.

| | Soil | Marine | Sheep rumen | Human gut | anaerobic digester sludge |
|---|---|---|---|---|---|
| # contigs (mean) | 1,073,856 | 30,955 | 69,658 | 18,830 | 35,334 |
| # contigs (max) | 1,073,856 | 52,389 | 69,658 | 21,318 | 40,430 |
| # contigs (min) | 1,073,856 | 12,547 | 69,658 | 16,703 | 30,043 |
| N50 (mean) | 35,862 | 44,466 | 290,707 | 175,316 | 108,352 |
| N50 (max) | 35,862 | 52,136 | 290,707 | 223,332 | 114,946 |
| N50 (min) | 35,862 | 33,975 | 290,707 | 135,807 | 95,956 |
| Total length (mean) | 24,703,618,963 | 805,772,635 | 4,513,827,413 | 1,036,218,753 | 1,485,366,378 |
| Total length (max) | 24,703,618,963 | 1,315,234,020 | 4,513,827,413 | 1,106,171,803 | 1,738,443,681 |
| Total length (min) | 24,703,618,963 | 306,681,005 | 4,513,827,413 | 958,496,208 | 1,263,125,584 |

**Supplementary Table 17** Overview of metaFlye-based Nanopore single-sample assemblies used for benchmarking.

| | Soil | Activated sludge |
|---|---|---|
| # contigs (mean) | 81,029 | 37,432 |
| # contigs (max) | 109,417 | 65,716 |
| # contigs (min) | 56,426 | 13,560 |
| N50 (mean) | 122,330 | 101,327 |
| N50 (max) | 194,707 | 191,848 |
| N50 (min) | 67,024 | 74,463 |
| Total length (mean) | 3,392,098,270 | 2,141,174,635 |
| Total length (max) | 3,772,507,743 | 3,566,969,444 |
| Total length (min) | 2,776,287,842 | 850,230,903 |

**Supplementary Table 18** Overview of Illumina single-sample assemblies used for benchmarking.

| | Marine | Activated sludge |
|---|---|---|
| # contigs (mean) | 436,809 | 324,059 |
| # contigs (max) | 506,460 | 419,437 |
| # contigs (min) | 326,241 | 227,648 |
| N50 (mean) | 2,590 | 2,289 |
| N50 (max) | 2,716 | 3,958 |
| N50 (min) | 2,526 | 2,000 |
| Total length (mean) | 1,026,002,035 | 702,196,567 |
| Total length (max) | 1,186,933,516 | 900,384,805 |
| Total length (min) | 787,174,817 | 492,893,842 |

**Supplementary Table 19** Overview of hybrid single-sample assemblies used for benchmarking.

| | Marine | Activated sludge |
|---|---|---|
| # contigs (mean) | 278,255 | 179,580 |
| # contigs (max) | 334,004 | 219,349 |
| # contigs (min) | 192,449 | 122,382 |
| N50 (mean) | 7,574 | 29,783 |
| N50 (max) | 11,056 | 43,883 |
| N50 (min) | 2,838 | 16,782 |
| Total length (mean) | 957,520,751 | 1,664,070,909 |
| Total length (max) | 1,227,382,210 | 2,251,057,464 |
| Total length (min) | 774,006,614 | 989,372,543 |
